# Natriuretic Peptide Augmentation Attenuates Renin Cell Hyperactivation and Afferent Arteriolar Hypertrophy During Long-Term Renin-Angiotensin System Inhibition

**DOI:** 10.64898/2026.07.21.739692

**Authors:** Takamitsu Shiiya, Hirofumi Watanabe, Ryo Aida, Chihiro Sakurazawa, Mizuki Honda, Yasuyuki Ohkawa, Shinya Oki, Tadashi Otsuka, Ryohei Kaseda, Shin Goto, Ichiei Narita, Suguru Yamamoto

## Abstract

**BACKGROUND:** Chronic renin-angiotensin system (RAS) inhibition activates renin cells and induces afferent arteriolar hypertrophy, a maladaptive vascular response that may contribute to nephrosclerosis-like renal injury. Although genetic or cell ablation approaches have shown that renin cells are required for this remodeling, no pharmacological strategy to restrain hyperactivated renin cells while preserving RAS inhibition benefits has been established. Natriuretic peptides (NPs) counteract RAS; however, whether NP signaling modulates renin cell activation and afferent arteriolar remodeling remains unclear.

**METHODS:** We examined direct effects of atrial natriuretic peptide (ANP) on As4.1 renin-producing cells using reverse transcription-quantitative PCR, ELISA, and RNA sequencing (RNA-seq). We established a mouse model of long-term RAS inhibition using valsartan, an angiotensin II receptor blocker (ARB), and compared it with sacubitril/valsartan, an angiotensin receptor-neprilysin inhibitor (ARNI). Valsartan was matched between the ARB and ARNI groups. Renin cell activation, afferent arteriolar remodeling, and renal injury were evaluated using biochemical assays, histology, immunostaining, single-nucleus RNA-seq, and region-specific photo-isolation chemistry RNA-seq of afferent arteriolar/juxtaglomerular regions.

**RESULTS:** ANP suppressed *Ren1* expression and renin secretion in As4.1 cells; this effect was attenuated by a natriuretic peptide receptor A antagonist. RNA-seq demonstrated that ANP induced receptor-dependent remodeling of renin cell gene programs. In mice, long-term ARB treatment induced renin cell hyperactivation, expansion of renin-positive juxtaglomerular regions, afferent arteriolar hypertrophy, renal dysfunction, tubular injury markers, and fibrosis. ARNI increased plasma ANP levels and attenuated these pathological changes, despite a comparable blood pressure reduction. Single-nucleus transcriptomics revealed attenuation of tubular injury-associated cellular states and altered renin cell-associated mesenchymal programs with ARNI. Region-specific transcriptomics further demonstrated distinct molecular states in afferent arteriolar/juxtaglomerular regions between ARB- and ARNI-treated kidneys. Integrated transcriptomic analysis suggested that NP signaling converges on vascular regulatory programs in hyperactivated renin-expressing cells.

**CONCLUSIONS:** NP signaling acts as a pharmacologically augmentable modulator of maladaptive renin cell activation. ARNI attenuates renin cell hyperactivation, afferent arteriolar hypertrophy, and renal injury during long-term RAS inhibition, suggesting that neprilysin inhibition may preserve RAS blockade benefits while limiting renin cell-driven renal vascular remodeling.

## BACKGROUND

Renin-angiotensin system (RAS) inhibitors, including angiotensin-converting enzyme (ACE) inhibitors and angiotensin II receptor blockers (ARBs), are widely used as first-line agents for hypertension treatment; they represent a cornerstone of cardiovascular medicine because of their antihypertensive and organ-protective effects, including the reduction of cardiovascular events^1–4^. The RAS is a fundamental regulator of physiological homeostasis, particularly blood pressure, extracellular fluid volume, and electrolyte balance^5^. Renin cells, located in the juxtaglomerular (JG) region of the afferent arteriole, serve as key regulatory nodes of this system by controlling renin synthesis and secretion, thereby maintaining systemic and renal hemodynamic homeostasis^6^. Renin cells are a highly specialized cell population that constitutes only a small fraction of renal cells^7^. During kidney development, renin-expressing cells are relatively abundant and broadly distributed throughout the renal vasculature. However, as development proceeds, their distribution becomes progressively restricted. In mature kidneys, they are predominantly localized to the JG region at the entrance of the glomerulus. Importantly, renin lineage cells retain remarkable plasticity. In response to threats to homeostasis, such as dehydration, hypotension, or sodium depletion, cells that previously expressed renin during development can reversibly reacquire a renin-expressing phenotype, thereby augmenting renin production and restoring physiological equilibrium^8–10^.

Chronic inhibition of the RAS is another potent stimulus that activates renin cells by interrupting feedback regulation. Previous studies using various animal models have shown that genetic or pharmacological suppression of the RAS induces marked hyperactivation and phenotypic remodeling of renin cells, resulting in hypertrophy of the renal afferent arteriole^11–15^. Moreover, long-term use of RAS inhibitors is associated with hypertrophy of afferent arterioles in the human kidney^12,16^. Renin cell hyperactivation and afferent arteriolar remodeling may reduce downstream renal blood flow, promote glomerular ischemia, and contribute to nephrosclerosis-like renal injury^12,17^. However, the molecular mechanisms underlying this process remain poorly understood, and there is no established pharmacological strategy to prevent maladaptive renin cell activation and afferent arteriolar hypertrophy while preserving the beneficial effects of RAS inhibition.

Natriuretic peptides (NPs) are endogenous counter-regulatory hormones that oppose the RAS in controlling body fluid balance, blood pressure, vascular tone, and neurohormonal activity^18,19^. Among the NP family members, atrial natriuretic peptide (ANP) promotes natriuresis and vasodilation, lowers blood pressure, suppresses sympathetic activity, and inhibits renin secretion and RAS activity^20–22^. These features suggest that NP signaling may counteract excessive renin cell activation. However, whether NPs act directly on renin cells and the molecular mechanisms that mediate their effects on renin expression and the renin cell phenotype remain unclear.

Sacubitril/valsartan, an angiotensin receptor-neprilysin inhibitor (ARNI), provides a clinically relevant approach for enhancing endogenous NP signaling during RAS inhibition. Sacubitril inhibits neprilysin, the enzyme responsible for NP degradation, whereas valsartan blocks angiotensin II type 1 receptor^23,24^. Thus, sacubitril/valsartan simultaneously augments NP signaling and suppresses the RAS. Clinically, sacubitril/valsartan is an established therapy for heart failure and has demonstrated meaningful blood pressure-lowering effects, including superiority over conventional ARB therapy in patients with hypertension^25–28^. In addition, emerging clinical evidence suggests that ARNI exerts cardiorenal protective effects in selected settings^27–29^. Nevertheless, it remains unknown whether ARNI induces afferent arteriolar hypertrophy in the same manner as conventional RAS inhibitors or whether neprilysin inhibition and NP augmentation mitigate renin cell hyperactivation and vascular remodeling during long-term RAS blockade.

In this study, we aimed to investigate whether NP augmentation modulates renin cell activation and afferent arteriolar remodeling during long-term RAS inhibition. Using cultured renin-producing cells, a novel mouse model of gradual long-term RAS inhibition, single-nucleus RNA sequencing (snRNA-seq), and region-specific photo-isolation chemistry RNA sequencing (PIC RNA-seq) of the afferent arteriolar/JG regions, we examined the effects of ANP and ARNI on hyperactivated renin cells. Our findings revealed that ANP directly suppresses renin expression through natriuretic peptide receptor A (NPRA) signaling and that ARNI attenuates renin cell hyperactivation, afferent arteriolar hypertrophy, and renal injury during chronic RAS inhibition. These findings identify NP signaling as a pharmacologically augmentable modulator of renin cell plasticity and suggest that ARNI may preserve the benefits of RAS inhibition while limiting maladaptive afferent arteriolar remodeling.

## RESULTS

### ANP suppresses renin expression and secretion in renin cells through NP receptor signaling

To determine whether NPs act directly on renin cells and suppress renin expression, we treated As4.1 cells, a mouse JG renin-producing cell line, with ANP (Figure 1A). Cells were treated with ANP twice at 12-h intervals and evaluated 24 h after the initial treatment. *Ren1* expression was significantly reduced by ANP in a dose-dependent manner (Figure 1B). Afterward, renin protein levels in the culture supernatants were measured using ELISA. Consistent with the mRNA results, renin concentrations in the culture supernatant significantly decreased after ANP treatment (Figure 1C).

**Figure 1.**
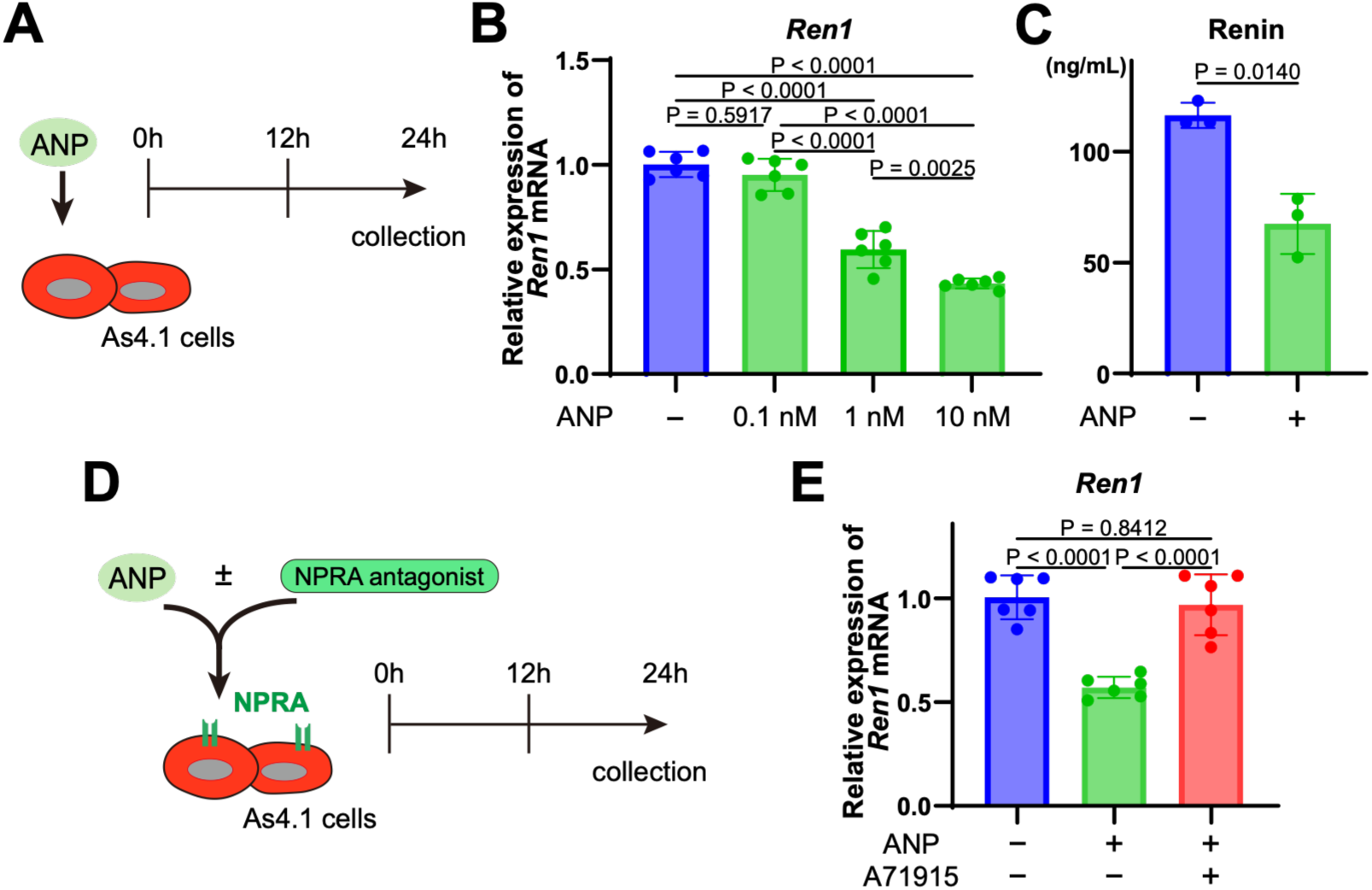
Atrial natriuretic peptide suppresses renin expression and secretion in cultured renin cells through natriuretic peptide receptor A signaling. **A**, Experimental scheme for atrial natriuretic peptide (ANP) treatment in As4.1 cells. ANP was added to the culture medium at 0 h and re-administered at 12 h, and the cells and culture supernatants were collected 24 h after the initial treatment. **B**, Relative *Ren1* mRNA expression 24 h after treatment with ANP at the indicated concentrations. *Ren1* expression was significantly suppressed by ANP in a dose-dependent manner (n = 6; one-way ANOVA followed by Tukey’s multiple comparison test). **C**, Renin concentrations in culture supernatants measured via ELISA after treatment with ANP (1 nM) or vehicle control (n = 3; Welch’s t-test). D, Experimental scheme of ANP treatment in the presence or absence of the natriuretic peptide receptor A antagonist A71915. **E**, Relative *Ren1* mRNA expression 24 h after treatment with ANP (1 nM) with or without A71915 (5 µM). A71915 significantly attenuated the inhibitory effect of ANP on *Ren1* expression (n = 6; one-way ANOVA, followed by Tukey’s multiple comparison test). Data are shown as mean ± SD. The cell culture data in B, C, and E are representative of three independent experiments.

To examine whether the suppressive effect of ANP on renin expression was mediated by NP receptor signaling, As4.1 cells were co-treated with ANP and the NPRA antagonist A71915 (Figure 1D). In the presence of A71915, the ANP-induced suppression of renin expression was significantly attenuated (Figure 1E, Figure S1). These findings indicate that ANP acts directly on renin cells through NPRA signaling to suppress renin expression.

### ANP induces NPRA-dependent transcriptional remodeling of renin cells

To elucidate the molecular mechanisms underlying the ANP-mediated suppression of renin gene expression, we performed transcriptome analysis. We performed RNA-seq analysis of As4.1 cells treated with vehicle, ANP, or ANP and the NPRA antagonist A71915. Pairwise differential expression analyses comparing control versus ANP-treated cells and ANP-treated versus ANP plus A71915-treated cells identified 2,413 and 1,831 differentially expressed genes (DEGs), respectively (Figure 2A, B). Consistent with the reverse transcription-quantitative polymerase chain reaction (RT-qPCR) results, *Ren1* expression was reduced by ANP treatment; this reduction was attenuated by NPRA antagonism. Principal component analysis (PCA) of the three groups after batch correction showed clear separation among the three groups. ANP-induced transcriptional changes were partially reversed toward the control state following NPRA inhibition (Figure 2C).

**Figure 2.**
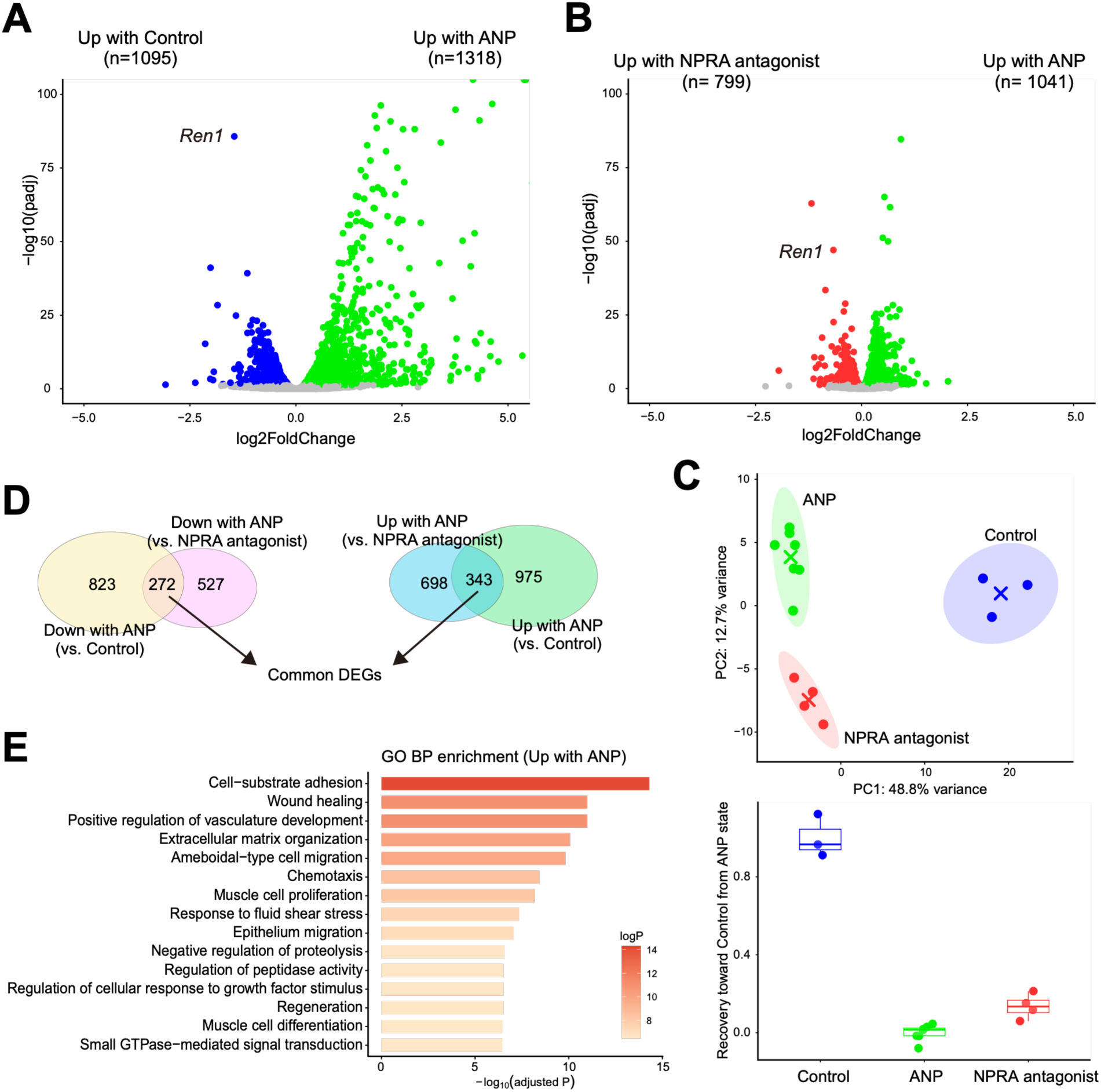
ANP induces natriuretic peptide receptor A-dependent transcriptional remodeling in renin cells. **A,** Volcano plot showing differentially expressed genes (DEGs) between the vehicle- and ANP-treated As4.1 cells. Significantly altered genes are highlighted (adjusted P < 0.05). **B,** Volcano plot showing DEGs between ANP-treated cells and cells treated with ANP plus the natriuretic peptide receptor A antagonist A71915. Significantly altered genes are highlighted (adjusted P < 0.05). **C,** Principal component analysis (PCA) of RNA-seq profiles from control, ANP-treated, and ANP plus A71915-treated cells after batch correction. ANP treatment induced a distinct transcriptional shift, which was partially reversed by A71915 treatment. **D,** Identification of 615 common DEGs with consistent directions of expression changes across two pairwise comparisons, including *Ren1*. **E,** Gene ontology (GO) Biological Process enrichment analysis of common DEGs upregulated by ANP in a natriuretic peptide receptor A-dependent manner.

To define the ANP-responsive, NPRA-dependent transcriptional program, we extracted genes that were altered by ANP and directionally reversed by A71915. This analysis identified 343 genes upregulated and 272 genes downregulated by ANP in an NPRA-dependent manner (Figure 2D). Gene ontology (GO) enrichment analysis of ANP-upregulated genes revealed the enrichment of biological processes related to cell adhesion, vascular development, extracellular matrix organization, cell migration, and muscle-associated functions (Figure 2E). In contrast, GO enrichment analysis of ANP-downregulated genes revealed the enrichment of terms related to cell division and cell cycle regulation (Figure S2).

Together, these findings indicate that ANP acts on renin cells through NPRA signaling to suppress renin expression and induce broad transcriptional remodeling toward a more contractile-like cellular state.

### Gradual long-term RAS inhibition induces renal dysfunction, which is attenuated by neprilysin inhibition

To investigate the renal consequences of gradual RAS inhibition and rigorously assess the effects of NPs on renin cell hyperactivation and afferent arteriolar hypertrophy, we established a novel chronic mouse model. Although mouse models of afferent arteriolar hypertrophy induced by long-term RAS inhibition have been previously reported^12^, we aimed to develop a model in which afferent arteriolar hypertrophy developed in the absence of overt renal injury during the early phase, thereby more closely recapitulating the gradual progression of nephrosclerosis-like renal injury in humans.

Eight-week-old C57BL/6 mice received once-daily oral gavage of either the ARB valsartan or the ARNI sacubitril/valsartan, were maintained on a low-salt diet with free access to water, and were followed longitudinally (Figure 3A). In the ARB group, afferent arteriolar hypertrophy progressively developed with prolonged treatment and significantly increased at 12 weeks (Figure S3A, S3B). However, no decline in renal function was observed in either treatment group for up to 12 weeks after treatment initiation (Figure S3C). These findings confirmed the successful establishment of a mouse model of gradual RAS inhibition-induced afferent arteriolar hypertrophy without overt renal injury during the early phase.

**Figure 3.**
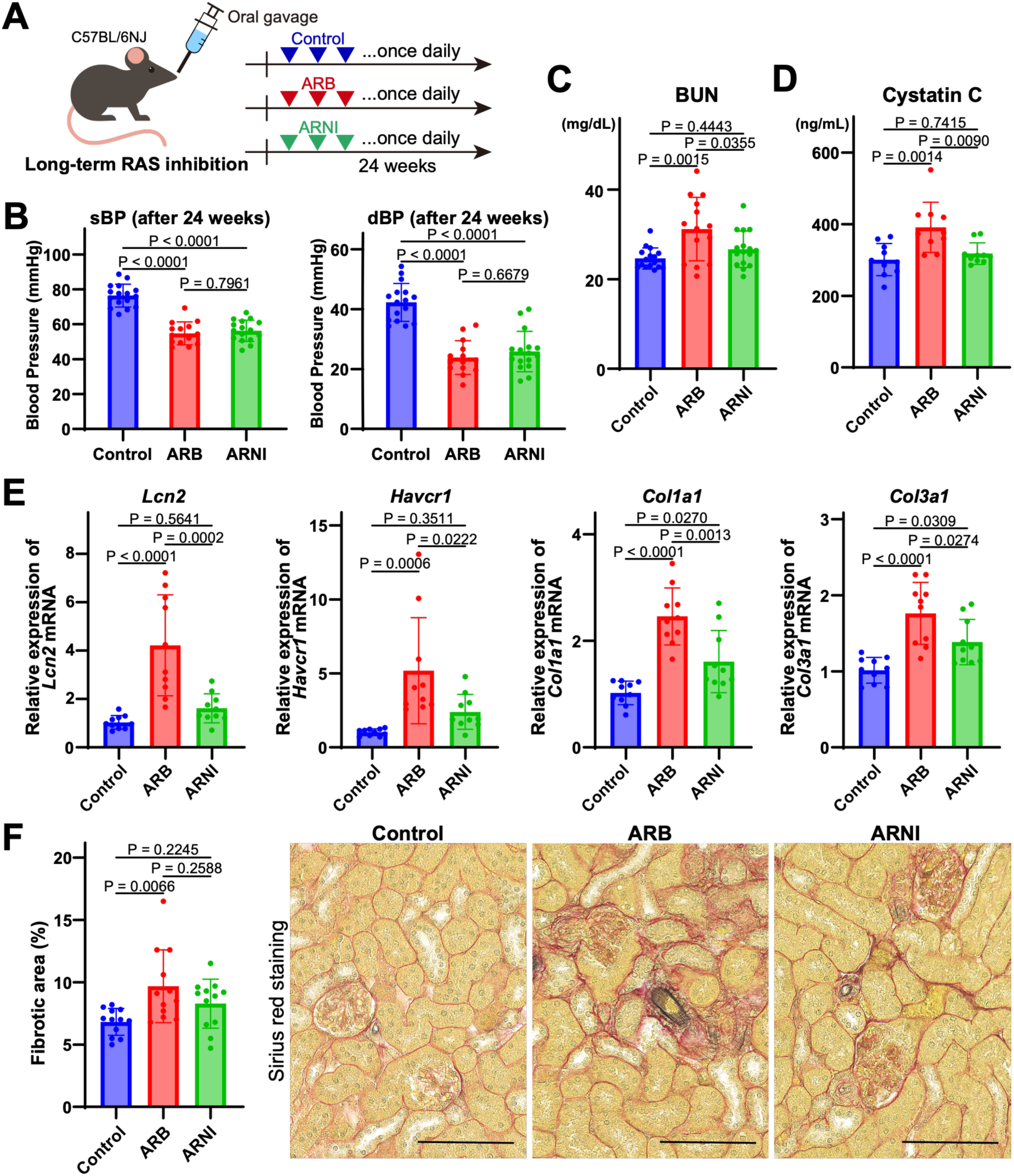
Long-term renin-angiotensin system inhibition induces renal dysfunction and fibrosis, which are attenuated by ARNI. **A,** Schematic overview of the long-term renin-angiotensin system (RAS) inhibition experiment in mice. Eight-week-old C57BL/6 mice received once-daily oral gavage of vehicle control (saline), the angiotensin II receptor blocker (ARB) valsartan, or the angiotensin receptor-neprilysin inhibitor (ARNI) sacubitril/valsartan. Blood and kidney samples were collected 24 weeks after treatment initiation. **B,** Systolic and diastolic blood pressure after 24 weeks of treatment, measured under isoflurane anesthesia using the tail-cuff method. After treatment, blood pressure was significantly lower in both the ARB and ARNI groups than in the control group, with no significant difference between the ARB and ARNI groups (n = 14 for ARB; n = 16 for control and ARNI; one-way ANOVA followed by Tukey’s multiple comparison test). **C,** Serum blood urea nitrogen (BUN) levels after 24 weeks of treatment. The BUN levels were significantly higher in the ARB group (n = 14 for ARB, n = 16 for control and ARNI; one-way ANOVA followed by Tukey’s multiple comparison test). **D,** Plasma cystatin C levels measured using ELISA. Plasma cystatin C levels were significantly elevated in the ARB group but not in the ARNI group (n = 10; one-way ANOVA followed by Tukey’s multiple comparison test). **E,** Relative mRNA expression of renal injury and fibrosis markers in kidney tissues. Long-term ARB treatment significantly increased the expression of *Lcn2*, *Havcr1*, *Col1a1*, and *Col3a1*, whereas these increases were attenuated in the ARNI group compared to those in the ARB group (n = 10; one-way ANOVA followed by Tukey’s multiple comparison test). **F,** Representative Sirius red-stained renal cortical sections and quantification of the fibrotic area. The ARB group showed a significant increase in renal fibrosis, whereas no significant increase was observed in the ARNI group. Scale bar, 100 µm. sBP, systolic blood pressure; dBP, diastolic blood pressure.

Next, we extended the drug administration to 24 weeks and performed detailed phenotypic analyses to evaluate the effects of NP augmentation on RAS inhibition. To isolate the contribution of neprilysin inhibition, mice in the ARB and ARNI groups received equivalent doses of valsartan, either as valsartan alone (30 mg/kg) or sacubitril/valsartan (60 mg/kg), once daily via oral gavage. This allowed us to assess the effects attributable to the presence or absence of sacubitril. Blood pressure measured under anesthesia at 24 weeks was significantly lower in both RAS inhibitor-treated groups than in the controls, with no significant differences between the ARB and ARNI groups (Figure 3B). Notably, renal function differed between the two treatment groups. In the ARB group, both BUN and plasma cystatin C levels were significantly elevated, whereas in the ARNI group, neither parameter differed significantly from that in the control group (Figure 3C, 3D). Similar differences were observed in intrarenal injury markers. The mRNA expression levels of the renal injury markers *Lcn2* (NGAL) and *Havcr1* (KIM-1) were significantly higher in the kidneys of the ARB group than in those of the control group, whereas no significant increase was observed in the ARNI group. In addition, the fibrosis markers *Col1a1* and *Col3a1* were upregulated in both the ARB and ARNI groups relative to the controls, although the magnitude of induction was attenuated in the ARNI group (Figure 3E). Consistent with these findings, Sirius red staining revealed significant renal fibrosis in the ARB group, whereas no significant difference was observed between the ARNI and control groups (Figure 3F).

Collectively, these results indicate that, in our model, gradual ARB-induced afferent arteriolar hypertrophy leads to renal dysfunction after prolonged treatment, and this process is mitigated by neprilysin inhibition.

### NPs attenuate hyperactivation of renin cells and afferent arteriolar hypertrophy

Next, we sought to elucidate why ARNI, compared to ARB, provided greater protection against renal dysfunction associated with long-term RAS inhibition. We confirmed elevated plasma ANP levels in the ARNI group (Figure 4A), indicating that NP signaling was enhanced in this group. To determine RAS inhibitor-induced hyperactivation of renin cells, we first examined *Ren1* mRNA expression in whole kidney samples. *Ren1* expression in the ARB group was significantly increased to approximately 20–30 times that in the control group (Figure 4B). Additionally, *Ren1* expression was elevated in the ARNI group; however, the magnitude of this increase was less pronounced than that observed in the ARB group. Consistently, plasma renin concentrations measured via ELISA were significantly elevated in both the ARB and ARNI groups compared with those in the control group, although the increase was attenuated in the ARNI group (Figure 4C).

**Figure 4.**
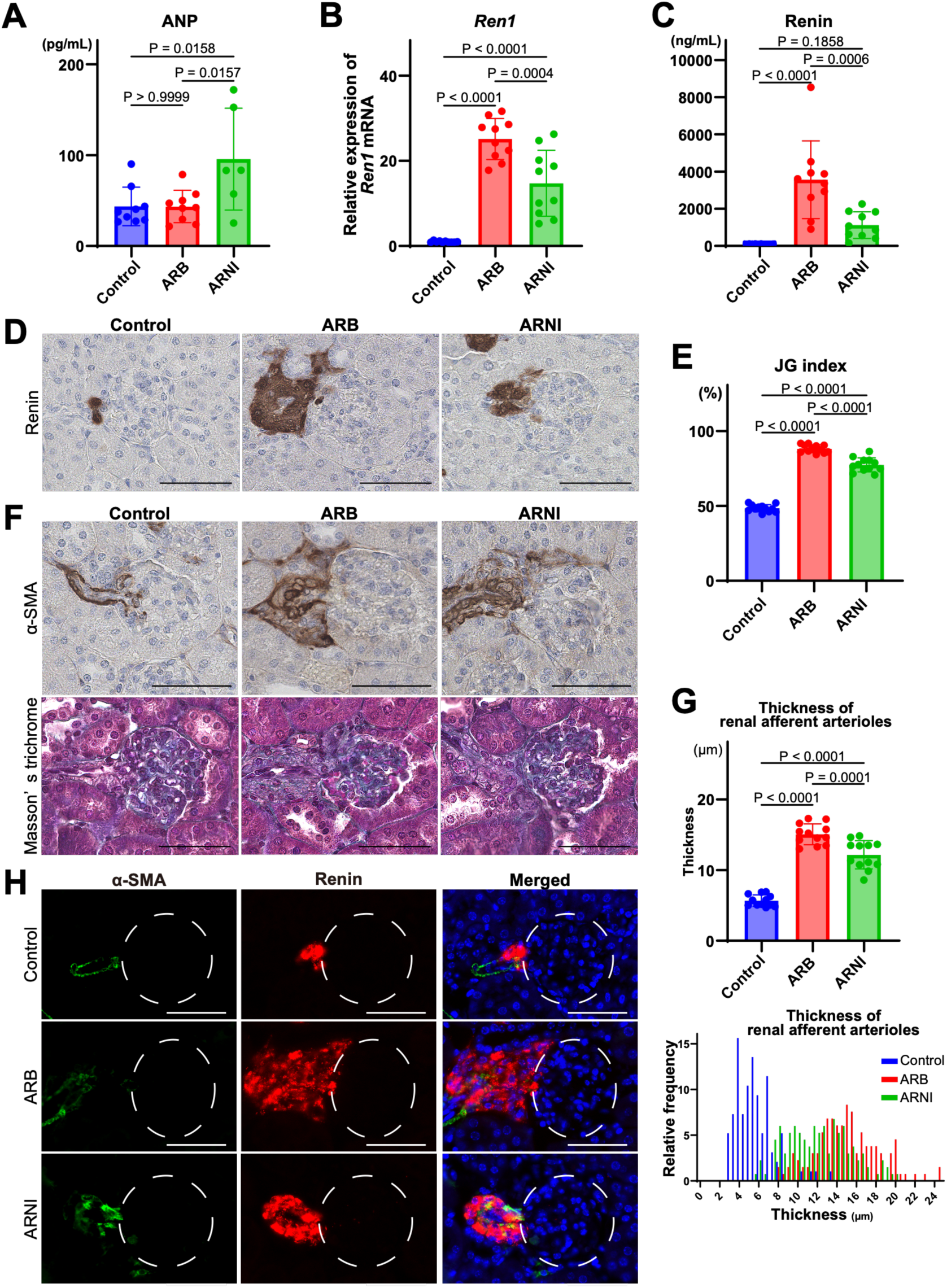
ARNI attenuates renin cell hyperactivation and afferent arteriolar hypertrophy induced by long-term RAS inhibition. **A,** Plasma atrial natriuretic peptide (ANP) levels measured using ELISA. Plasma ANP levels significantly increased only in the ARNI group (n = 9 for control and ARB; n = 6 for ARNI; one-way ANOVA followed by Tukey’s multiple comparison test). **B,** Relative Ren1 mRNA expression in kidney tissues. Long-term ARB treatment markedly increased Ren1 expression, whereas this upregulation was significantly attenuated in the ARNI group compared to that in the ARB group (n = 10; one-way ANOVA followed by Tukey’s multiple comparison test). **C,** Plasma renin concentrations measured using ELISA. Plasma renin levels were significantly elevated in both the ARB and ARNI groups compared to the control group, although the increase was significantly attenuated in the ARNI group compared to the ARB group. (n = 10; one-way ANOVA followed by Tukey’s multiple comparison test). **D,** Representative renin immunohistochemistry in mouse kidneys after 24 weeks of treatment. Scale bar, 50 µm. **E,** quantification of the juxtaglomerular (JG) index. The JG index, defined as the percentage of renin-positive JG apparatuses among the total glomeruli, was significantly increased by long-term RAS inhibition; this increase was attenuated in the ARNI group compared to that in the ARB group (n = 12, one-way ANOVA followed by Tukey’s multiple comparison test). **F,** Representative α-smooth muscle actin (α-SMA) immunohistochemistry and Masson’s trichrome staining of renal tissues. Scale bar, 50 µm. **G,** Quantification of afferent arteriolar wall thickness. Wall thickness was calculated as half the difference between the outer and inner diameters of the afferent arteriole. Long-term RAS inhibition significantly increased afferent arteriolar wall thickness, whereas this increase was attenuated in the ARNI group compared to the ARB group (n = 12; one-way ANOVA followed by Tukey’s multiple comparison test). The lower panel shows the relative frequency distribution of afferent arteriolar wall thickness measurements in each treatment group (control, blue; ARB, red; ARNI, green). **H,** Immunofluorescence staining for α-SMA and renin in afferent arteriolar/JG regions. The left panel shows α-SMA, the middle panel shows renin, and the right panel shows the merged image. Scale bar, 50 µm.

Furthermore, we performed renin immunohistochemistry on paraffin-embedded kidney sections and quantified the proportion of renin-positive glomeruli (JG index) (Figure 4D). The JG index was approximately 50% in the control group and was markedly increased to approximately 90% in the ARB group. In contrast, the JG index was significantly lower in the ARNI group than in the ARB group (Figure 4E). Measurement of the afferent arteriolar wall thickness in Masson’s trichrome-stained kidney sections revealed that wall thickening in the ARB group was approximately three-fold greater than that in the control group, whereas this increase was attenuated in the ARNI group (Figure 4F, 4G). Consistent with these findings, immunofluorescence analysis confirmed that ARNI markedly suppressed renin cell-associated structural changes in the afferent arteriole induced by long-term ARB treatment (Figure 4H).

Taken together, these results indicate that ARNI increases natriuretic peptide levels, thereby suppressing renin cell hyperactivation and afferent arteriolar hypertrophy induced by long-term RAS inhibition.

### Single-nucleus transcriptomics reveals attenuation of renal injuries by ARNI during long-term RAS inhibition

To capture the cell type-specific changes induced by ARB or ARNI treatment, we performed snRNA-seq using mouse kidneys 24 weeks after drug administration. Nuclei were isolated from three kidney cortices per group (control, ARB, and ARNI). After sequencing and quality filtering, we obtained 5,844, 1,442, and 1,200 nuclei from the control, ARB, and ARNI groups, respectively. We integrated all datasets, visualized the transcriptomic landscape using uniform manifold approximation and projection (UMAP), and identified and assigned 10 distinct cell populations based on canonical marker gene expression (Figure 5A).

**Figure 5.**
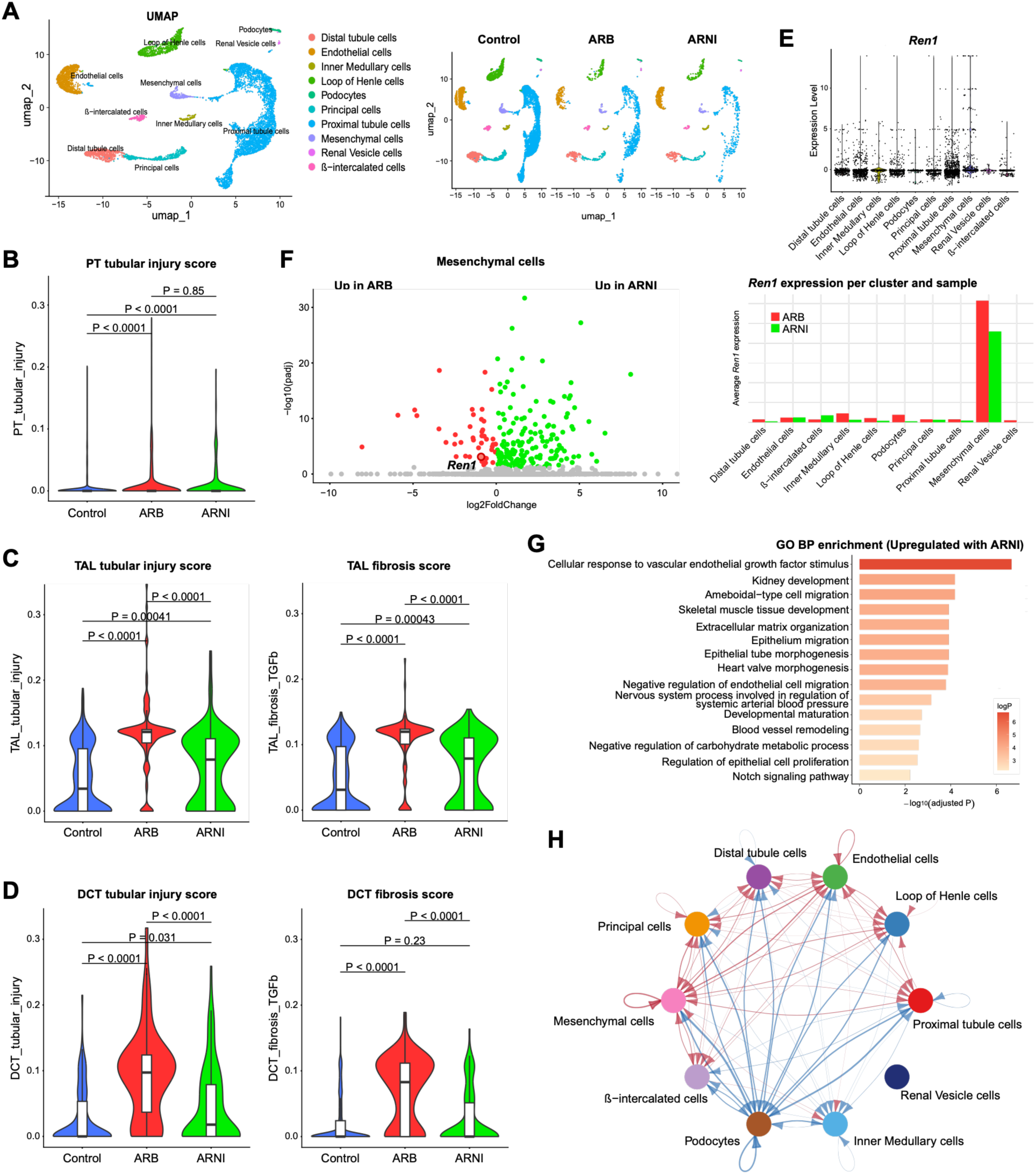
Single-nucleus RNA sequencing reveals ARNI-associated attenuation of tubular injury states and remodeling of renin cell-associated mesenchymal programs during long-term RAS inhibition. **A,** Single-nucleus RNA sequencing (snRNA-seq) was performed on the mouse renal cortices after 24 weeks of treatment. Integrated nuclei from the control, ARB, and ARNI groups were clustered and annotated using ScType. Uniform manifold approximation and projection (UMAP) visualization is shown by cell type annotation (left) and treatment group (right). Control, blue; ARB, red; ARNI, green. **B,** Reanalysis of proximal tubular clusters. Tubular injury activity was quantified using UCell signature scores based on a manually curated gene set comprising *Havcr1*, *Krt20*, *Krt8*, *Krt18*, *Krt19*, *Sox9*, *Clu*, *Vim*, *Spp1*, and *Egr1*. UCell scores were visualized using violin plots with overlaid boxplots. Pairwise comparisons were performed using Wilcoxon rank-sum tests. **C**, UCell-based injury and fibrosis scores in thick ascending limb (TAL) clusters. Tubular injury scores were calculated using a gene set consisting of *Lcn2*, *Havcr1*, *Krt8*, *Krt18*, *Spp1*, *Clu*, *Vim*, and *Sox9*. Fibrosis-associated scores were calculated using a gene set comprising *Tgfbi*, *Serpine1*, *Thbs1*, *Fn1*, *Col1a1*, *Col3a1*, *Tagln*, and *Spp1*. **D**, UCell-based injury and fibrosis scores in the distal convoluted tubule (DCT) clusters. Tubular injury and fibrosis-associated scores were calculated using the same gene sets as those in **C**. E, *Ren1* expression in the snRNA-seq dataset. Violin plots of *Ren1* expression are shown on the left, and bar plots of the average *Ren1* expression across cell clusters and treatment groups are shown on the right. The mesenchymal cell cluster showed the highest *Ren1* expression, which was higher in the ARB group than in the ARNI group. **F,** Volcano plot showing differentially expressed genes between the ARB and ARNI groups within the mesenchymal cell cluster. Significantly altered genes are highlighted in red, with an adjusted P value < 0.05. *Ren1* expression was also higher in the ARB group. **G,** Gene ontology (GO) Biological Process enrichment analysis of differentially expressed genes in the mesenchymal cell cluster between the ARB and ARNI groups. Enriched terms associated with the genes upregulated in the ARNI group are shown. **H,** CellChat analysis comparing the inferred cell-cell communication networks between the ARB and ARNI groups. Changes in the number of interactions between cell clusters are indicated by arrows; interactions that increased in the ARB group are shown in blue, and those that increased in the ARNI group are shown in red.

Compared to the control group, the ARB group showed transcriptional features consistent with renal injury. In proximal tubular cells, GO enrichment analysis of DEGs revealed that several immune response-related terms were among the most enriched biological processes in the ARB group compared to the control group. By contrast, these terms were less prominently enriched in the ARNI group (Figure S4). Next, we quantified predefined tubular injury gene signatures using UCell^30^. In the proximal tubular clusters, the tubular injury marker scores did not differ significantly between the ARB and ARNI groups (Figure 5B). However, in downstream tubular segments, including the loop of Henle and distal tubule clusters, the injury and fibrosis scores increased in the ARB group, whereas these increases were attenuated in the ARNI group (Figure 5C, D).

Together, these findings suggest that, at single-nucleus resolution, ARNI is associated with attenuation of tubular injury-associated cellular states induced by long-term RAS inhibition.

### ARNI alters renin cell-associated mesenchymal programs and endothelial interactions during long-term RAS inhibition

Renin cells, identified by *Ren1* expression, were located within the mesenchymal cell cluster via snRNA-seq (Figure 5E). The average *Ren1* expression per cell within the mesenchymal cell cluster was higher in the ARB group than that in the ARNI group (Figure 5E). Afterward, we performed differential gene expression analysis within the mesenchymal cell cluster between the ARB and ARNI groups. This analysis identified 160 and 38 upregulated genes in the ARNI and ARB groups, respectively, with *Ren1* showing the highest expression in the ARB group (Figure 5F). These findings indicate that ARB and ARNI treatment induce distinct transcriptional states within the mesenchymal cell cluster containing renin-expressing cells.

To explore the molecular programs altered by ARNI treatment in renin cell-associated mesenchymal populations, we performed GO enrichment analysis of DEGs between the ARB and ARNI groups within the mesenchymal cell cluster identified via snRNA-seq. Genes upregulated in the ARNI group were significantly enriched for biological processes related to cardiovascular development, endothelial maturation and stabilization, and vascular remodeling. These included the regulation of cellular response to vascular endothelial growth factor stimulus, striated muscle tissue development, heart valve morphogenesis, negative regulation of endothelial cell migration, blood vessel remodeling, negative regulation of cellular response to vascular endothelial growth factor stimulus, regulation of vascular permeability, and regulation of muscle system processes (Figure 5G). In contrast, genes that were relatively downregulated in the ARNI group were enriched for cytokine-mediated signaling pathways (Figure S5A), suggesting the attenuation of inflammatory or neuroimmune-associated responses^31–33^. Consistent with this possibility, the expression of *Ngf*, a neurotrophic factor induced by RAS inhibition^15^, was less pronounced in the ARNI group (Figure S5B).

Next, we performed CellChat analysis^34^ to assess whether communication between mesenchymal cell clusters containing renin cells and other renal cell populations was altered by ARNI treatment. This analysis suggested increased cell-cell interactions between the mesenchymal and endothelial cell clusters in the ARNI group (Figure 5H).

Together, these findings suggest that ARNI modifies renin cell-associated mesenchymal programs and enhances mesenchymal-endothelial interactions during long-term RAS inhibition.

### JG region-specific spatial transcriptomic analysis reveals the effects of NP augmentation on hyperactivated renin cells

To further analyze the characteristics of the JG region and hypertrophied afferent arterioles, we performed PIC RNA-seq using mouse kidney tissues after 24 weeks of treatment. PIC RNA-seq enables region-specific transcriptomic profiling of tissue areas that are selectively targeted by light irradiation^35,36^. We stained kidney tissues from mice treated with either ARB or ARNI for 24 weeks with α-SMA to identify the afferent arteriolar/JG regions and performed targeted light irradiation. Light irradiation was applied to 20–30 afferent arterioles of four samples each from the ARB and ARNI groups, followed by region-specific transcriptome analysis of the irradiated areas (Figure 6A, B).

**Figure 6.**
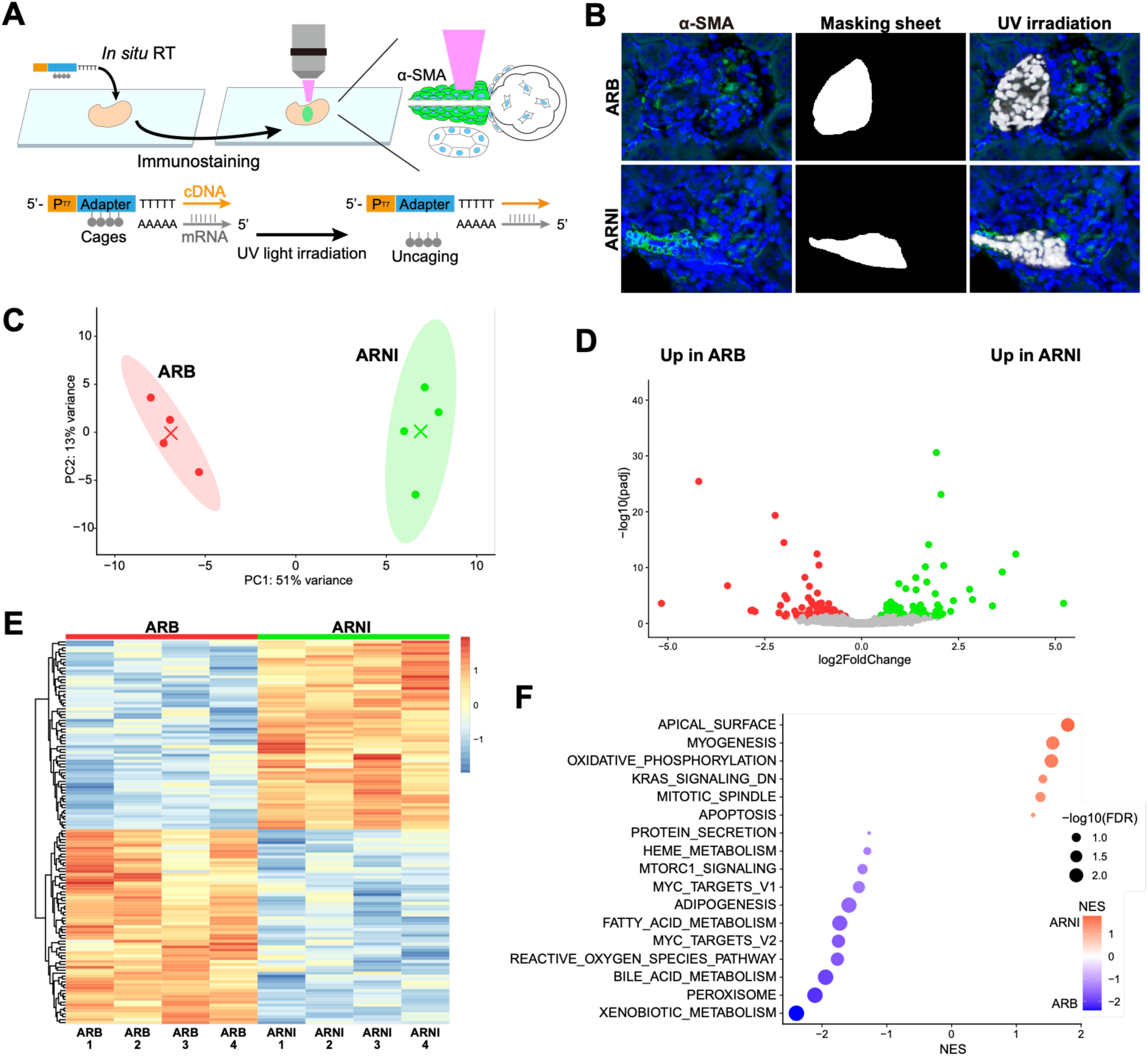
Region-specific PIC RNA-seq reveals distinct transcriptional states in afferent arteriolar/JG regions in ARNI-treated kidneys compared to ARB-treated kidneys. **A,** Schematic overview of photo-isolation chemistry (PIC) RNA-seq. Targeted light irradiation was applied to α-SMA-positive afferent arteriolar/juxtaglomerular (JG) regions in kidney sections from the ARB and ARNI groups after 24 weeks of treatment (n = 4 per group). **B,** Representative image of a kidney section used for targeted irradiation. Regions of interest were defined on α-SMA-stained sections as indicated by the masking sheet, and light irradiation was selectively applied to the white regions. **C,** Principal component analysis (PCA) of PIC RNA-seq profiles from the ARB and ARNI groups. **D,** Volcano plot showing differentially expressed genes between the ARB and ARNI groups in the irradiated afferent arteriolar/JG regions. Significantly altered genes are highlighted (adjusted P < 0.05). **E,** Heat map of differentially expressed genes between the ARB and ARNI groups. Region-specific gene expression profiles differed between the two groups within the afferent arteriolar/JG regions. **F,** Dot plot of hallmark gene set enrichment analysis (GSEA). Gene sets enriched in the ARNI group are shown in red, and those enriched in the ARB group are shown in blue. Circle size indicates −log10(FDR).

PCA revealed that gene expression profiles of individual samples exhibited distinct patterns according to the treatment group (Figure 6C). Differential expression analysis identified 67 upregulated genes in the ARNI group and 67 upregulated genes in the ARB group (Figure 6D). Visualization of the DEGs between the ARB and ARNI groups using a heat map showed that the two groups exhibited distinct expression patterns (Figure 6E).

To gain biological insight into treatment-dependent transcriptional differences in the afferent arteriolar/JG regions, we performed a hallmark gene set enrichment analysis. Gene sets related to the apical surface, myogenesis, and oxidative phosphorylation were highly enriched in the ARNI group, whereas gene sets related to xenobiotic metabolism, peroxisomes, bile acid metabolism, reactive oxygen species pathway, MYC targets, and fatty acid metabolism were predominantly enriched in the ARB group (Figure 6F).

Together, these findings indicate that ARNI-mediated NP augmentation alters the molecular state of hyperactivated renin cell-associated afferent arteriolar/JG regions.

### NP signaling may preserve afferent arteriolar vascular identity by remodeling hyperactivated renin cells

To explore the transcriptional programs through which NP signaling modulates renin-expressing cells, we integrated the snRNA-seq dataset with the RNA-seq data from ANP-treated As4.1 cells. Specifically, we searched for DEGs showing concordant directionality between the mesenchymal cell cluster containing *Ren1*-expressing cells in snRNA-seq and ANP-responsive genes in As4.1 cells. This approach identified 17 core genes associated with NP-mediated renin cell remodeling, including *Ren1*, whose direction of change was consistent across datasets (Figure 7A). We performed GO Biological Process enrichment analysis of these genes. The enriched terms included pathways related to vascular endothelial growth factor (VEGF) signaling, MAPK signaling, cellular proliferation, and differentiation (Figure 7B). A GO term-gene network further showed that these pathways were connected by multiple shared genes, suggesting coordinated regulation of vascular and growth factor-responsive programs (Figure 7C).

**Figure 7.**
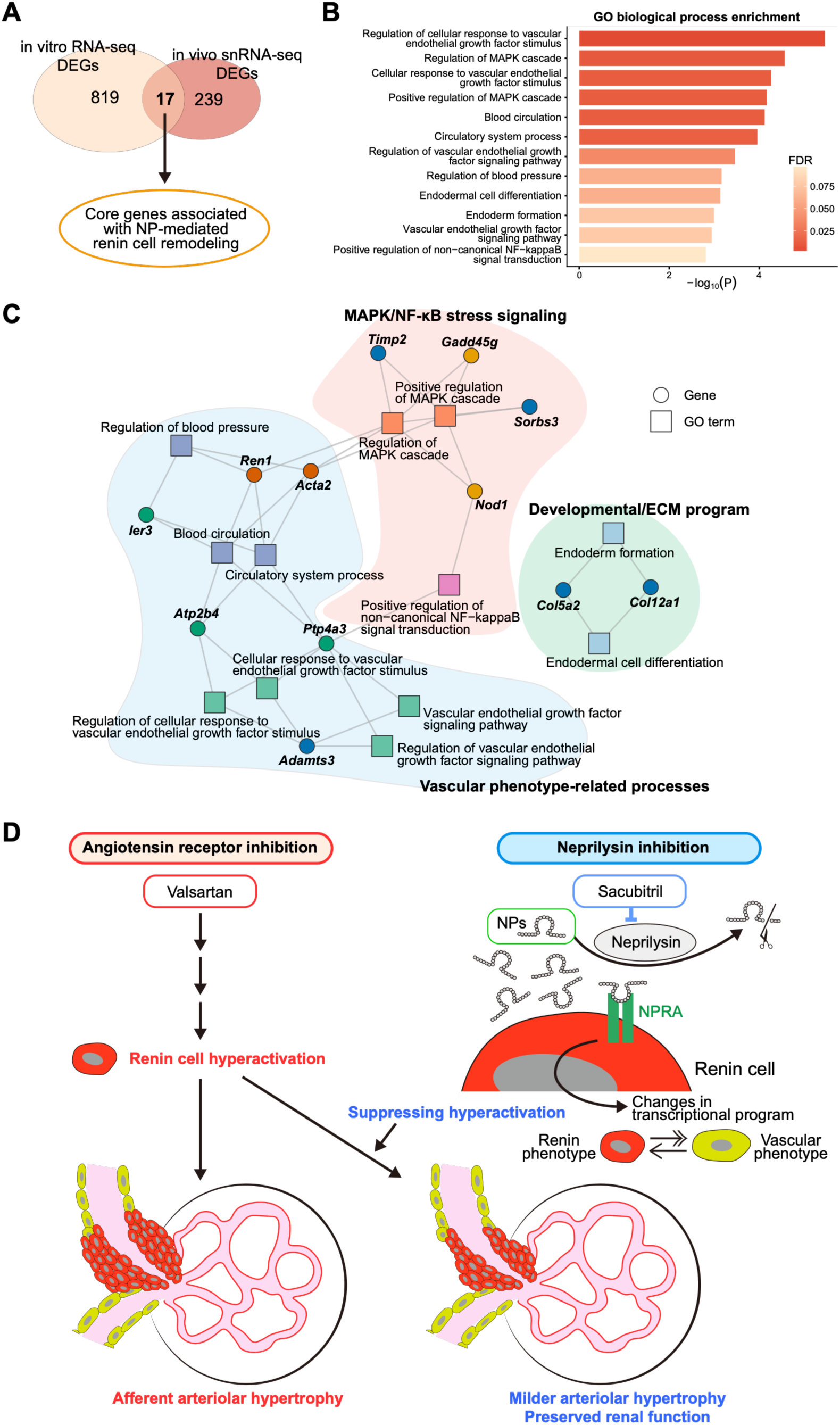
Integrated transcriptomic analysis identifies NP-responsive vascular regulatory programs in hyperactivated renin cells. **A,** Identification of shared candidate genes linking NP signaling to renin cell remodeling. Differentially expressed genes were compared between ANP-treated As4.1 cells and the mesenchymal cell cluster containing *Ren1*-expressing cells in the snRNA-seq dataset. Genes showing concordant directions of change in both datasets were selected using an adjusted P value < 0.1 as an exploratory threshold. This analysis identified 17 shared candidate genes, including *Ren1*. **B**, Gene ontology (GO) Biological Process enrichment analysis of 17 shared candidate genes. Enriched terms with FDR < 0.1 are shown. **C**, GO term-gene network plot showing the relationships between enriched biological processes and their associated genes. GO terms are shown as squares, genes as circles, and connecting lines indicate term-gene associations. **D**, Proposed model. Long-term RAS inhibition induces renin cell hyperactivation and afferent arteriolar hypertrophy. ARNI augments NP signaling by inhibiting neprilysin-mediated NP degradation. NP signaling suppresses renin cell hyperactivation and promotes vascular regulatory programs, thereby attenuating maladaptive phenotypic remodeling and preserving afferent arteriolar vascular identity.

Collectively, these exploratory analyses suggest that NP signaling converges with vascular regulatory programs in hyperactivated renin cells. During long-term RAS inhibition, NP augmentation may attenuate maladaptive renin cell remodeling and help preserve or partially restore the vascular identity of afferent arteriolar cells that have reacquired a renin-expressing phenotype (Figure 7D).

## DISCUSSION

This study demonstrated that NP signaling suppresses maladaptive renin cell activation and afferent arteriolar remodeling during long-term RAS inhibition. Using cultured renin-producing cells, we found that ANP directly suppressed *Ren1* expression and renin secretion via NPRA signaling. In mice, gradual long-term ARB treatment induced renin cell hyperactivation, afferent arteriolar hypertrophy, renal dysfunction, tubular injury marker induction, and fibrosis. Importantly, these pathological changes were attenuated by sacubitril/valsartan despite an equivalent valsartan component and a comparable blood pressure reduction. Single-nucleus and region-specific transcriptomic analyses further suggested that NP augmentation modified renin cell-associated mesenchymal and afferent arteriolar/JG transcriptional programs. Together, these findings identify NP signaling as a pharmacologically augmentable pathway that restrains excessive renin cell activation and protects against renin cell-associated renal vascular remodeling.

RAS inhibitors are indispensable therapeutic agents owing to their well-established antihypertensive, cardioprotective, and renoprotective effects, particularly in patients with hypertension, heart failure, diabetes, and proteinuric chronic kidney disease (CKD)^1,2,37,38^. However, chronic RAS inhibition inevitably activates renin cells by interrupting feedback regulation. In mature kidneys, renin-producing cells are primarily localized in the JG portion of the afferent arteriole. Under homeostatic stress, such as hypotension, dehydration, or chronic RAS inhibition, renin-lineage cells and afferent arteriolar smooth muscle cells reacquire a renin-expressing phenotype^6,10,39^. This response is essential for systemic homeostasis; however, when sustained, it may become maladaptive.

Previous studies have reported that long-term genetic or pharmacological suppression of the RAS induces hyperactivation of renin cells, recruitment of renin-expressing cells along the afferent arteriole, and concentric afferent arteriolar hypertrophy in mice and humans^12^. Our study extends these observations by showing that, even when overt renal injury is absent during the early phase, prolonged gradual RAS inhibition can eventually result in renal dysfunction, tubular injury marker induction, and fibrosis. This finding may be clinically relevant to nephrosclerosis-like kidney diseases. Nephrosclerosis is histologically heterogeneous; however, vascular narrowing, arteriolar wall thickening, and ischemic glomerular injury are important components of its pathology^40–42^. The rationale for RAS inhibition in patients with CKD and proteinuria is strong and supported by clinical trials and guidelines^37,38^. In contrast, the kidney-specific benefit of RAS inhibition in patients without albuminuria is less firmly established. Current CKD blood pressure guidelines primarily recommend ACE inhibitors or ARBs for patients with moderately to severely increased albuminuria, whereas their use in patients without albuminuria is framed more cautiously^37^. Our data did not contradict the use of RAS inhibitors. Rather, they raised the possibility that, in selected settings such as long-term exposure in kidneys susceptible to ischemic arteriolar remodeling, excessive renin cell activation may represent an underrecognized component of renal injury. Therefore, the chronic mouse model developed in this study may provide a useful experimental platform for investigating slowly progressive nephrosclerosis-like arteriolar remodeling.

A major conceptual advance of this study was the identification of a pharmacological approach capable of attenuating hyperactivated renin cells and afferent arteriolar hypertrophy. To our knowledge, previous experimental strategies to suppress renin cell hyperactivation or afferent arteriolar remodeling relied primarily on genetic or cell ablation approaches, such as conditional deletion of *Itgb1* in renin cells, which induces the loss of renin cells through apoptosis^12^, or diphtheria toxin-mediated ablation of renin cells^43^. These models are essential for demonstrating the requirement of renin cells in the development of afferent arteriolar hypertrophy. However, they do not represent clinically applicable interventions. In contrast, our findings showed that NP signaling, an endogenous counter-regulatory hormonal system, suppresses renin cell activation without eliminating renin cells. This distinction is important: rather than ablating the renin cell compartment, NP augmentation appears to modulate the pathological state of renin-lineage cells and redirect their transcriptional program toward a less activated, more vascular-like phenotype.

NPs and the RAS exert opposing physiological effects in the regulation of blood pressure, sodium balance, vascular tone, and neurohormonal activity^18,19^. ANP inhibits renin release from JG cells through a cGMP-dependent process^22^. In addition, RAS activation has been reported in mice lacking *Npr1*^44,45^, a gene encoding NPRA^20,46^. Consistent with these observations, ANP suppressed both *Ren1* mRNA expression and renin secretion in As4.1 cells in our study, and this effect was attenuated by the NPRA antagonist A71915^47,48^. These findings indicate that ANP acts directly on renin-producing cells, at least in part, through NPRA signaling. Furthermore, RNA-seq showed that ANP not only suppressed *Ren1* expression but also broadly remodeled the renin cell transcriptome. ANP-responsive genes were enriched for vascular development, cell adhesion, extracellular matrix organization, cell migration, and muscle-associated biological processes, whereas the downregulated genes were enriched for cell-cycle-related pathways. These findings suggest that ANP shifts renin cells away from an activated renin-producing state toward a more vascular, adhesive, and contractile-like transcriptional state.

In vivo ARNI experiments provide important translational extensions of these findings. Because ANP has a very short biological half-life^23,49^, direct chronic ANP administration is not an ideal strategy for long-term experiments. Sacubitril/valsartan allowed us to augment endogenous NP signaling during RAS inhibition. In our model, both ARB and ARNI lowered blood pressure to a similar extent, and the valsartan component was matched between the groups. However, ARNI attenuated the increase in renal *Ren1* expression, plasma renin levels, renin-positive JG regions, and afferent arteriolar wall thickness. These effects were accompanied by the preservation of renal function and the reduction of tubular injury and fibrosis. Thus, the protective effect of ARNI cannot be explained simply by a greater blood pressure reduction or a weaker RAS blockade. Rather, the data support the interpretation that neprilysin inhibition and NP augmentation directly modulate the intrarenal renin cell response during chronic RAS inhibition.

ARNI may preserve the systemic benefits of RAS inhibition while attenuating the maladaptive activation of the intrarenal renin cell compartment. Sacubitril/valsartan is an established therapy for heart failure, particularly heart failure with reduced ejection fraction^27^. It has clinically meaningful blood pressure-lowering effects, including favorable effects on central hemodynamic parameters, compared with conventional ARB therapy^50,51^. In addition to these cardiovascular and antihypertensive effects, clinical analyses in patients with heart failure have suggested that sacubitril/valsartan may slow eGFR decline and reduce serious renal outcomes compared to conventional RAS inhibition^28,29,52^. However, kidney-specific benefits outside the heart failure setting remain less certain; in patients with CKD, sacubitril/valsartan showed similar effects on measured GFR and albuminuria compared with irbesartan over 12 months, although it provided additional reductions in blood pressure and cardiac biomarkers^25^. Thus, the renal effects of ARNI are likely context dependent. Our findings provide a potential mechanistic explanation for how neprilysin inhibition may confer renal protection in selected settings: NP augmentation may directly modulate renin cell activation and afferent arteriolar remodeling beyond blood pressure lowering alone. Future clinical studies, particularly those incorporating kidney pathology or biomarkers of renin cell activation, are required to determine whether this mechanism is relevant in patients with hypertension, CKD, or nephrosclerosis-like kidney disease.

The snRNA-seq data further supported the protective effects of ARNI at the cellular level. Although the changes in the proximal tubular injury scores were modest, the downstream tubular segments, including the loop of Henle and distal tubular clusters, showed increased injury- and fibrosis-associated transcriptional signatures in the ARB group, which were attenuated in the ARNI group. This finding supports the hypothesis that afferent arteriolar hypertrophy and altered renal microvascular perfusion cause downstream tubular stress and injury over time. In the mesenchymal compartment containing *Ren1*-expressing cells, ARB and ARNI induced distinct transcriptional states. The ARNI group showed a lower average *Ren1* expression than did the ARB group, together with the enrichment of programs related to cardiovascular development, endothelial maturation, vascular remodeling, and muscle system processes. These results suggest that ARNI does not simply reduce renin expression quantitatively but may also alter the qualitative cellular state of renin cell-associated mesenchymal populations.

PIC RNA-seq provided complementary spatial information by focusing on α-SMA-positive afferent arteriolar/JG regions. This approach revealed a clear separation between the ARB- and ARNI-treated samples, indicating that neprilysin inhibition alters the molecular state of hyperactivated renin cell-associated vascular regions. Hallmark gene set analysis showed enrichment of myogenesis and apical surface-related pathways in the ARNI group, whereas several metabolic and stress-related pathways were enriched in the ARB group. Although PIC RNA-seq does not isolate pure renin cells, these data reinforce the idea that NP augmentation modifies the transcriptional landscape of the afferent arteriolar/JG niche during long-term RAS inhibition.

By integrating the snRNA-seq and ANP-treated As4.1 RNA-seq datasets, we identified a small set of candidate core genes that showed concordant changes across the in vivo and in vitro systems. These genes were enriched for biological processes related to VEGF signaling, MAPK signaling, cellular proliferation, and differentiation. Although exploratory, this integrated analysis suggests that NP signaling converges with vascular regulatory programs in hyperactivated renin-expressing cells. During chronic RAS inhibition, afferent arteriolar cells that reacquire renin expression may partially lose their original vascular identity and adopt a maladaptive endocrine remodeling phenotype. NP augmentation may counteract this process by preserving or partially restoring the vascular-associated cellular programs. This proposed mechanism provides a unifying explanation for the observed reduction in renin cell hyperactivation, afferent arteriolar hypertrophy, tubular injury, and fibrosis in the ARNI group.

This study had several limitations. First, although sacubitril/valsartan increased plasma ANP, and our in vitro data support a direct suppressive effect of ANP on renin cells, neprilysin has multiple substrates^53^. Therefore, the protective effects of sacubitril/valsartan in vivo cannot be attributed exclusively to ANP; other NPs or neprilysin substrates may also contribute. Second, the precise in vivo mechanism by which NP signaling modulates renin cell plasticity remains unclear. In particular, we did not directly investigate whether NPRA signaling in renin-lineage cells is required for the attenuation of renin cell hyperactivation and afferent arteriolar hypertrophy. Third, whether long-term ARNI therapy prevents afferent arteriolar hypertrophy or nephrosclerosis-like injury in human kidneys remains unknown. Future studies using human kidney tissue, pathology-based cohorts, or longitudinal clinical data are required to determine the clinical relevance of this mechanism.

In conclusion, this study identified NP signaling as a novel endogenous brake for maladaptive renin cell activation. We showed that ANP directly suppresses renin expression and secretion through NP receptor signaling, and that ARNI attenuates renin cell hyperactivation, afferent arteriolar hypertrophy, and renal dysfunction during long-term RAS inhibition. Unlike previous genetic- or ablation-based approaches, NP augmentation represents a pharmacologically accessible strategy for modulating renin cell plasticity without eliminating renin cells. These findings suggest that targeting the phenotypic state of renin-lineage cells may be a promising approach to preserve the cardiovascular and renal benefits of RAS inhibition, while preventing chronic afferent arteriolar remodeling and nephrosclerosis-like renal injury.

## METHODS

### Cell culture

The mouse renin cell line As4.1(CRL2193)^54^ was used. Cells were purchased from the American Type Culture Collection (Manassas, VA, USA). The cells were maintained in high-glucose Dulbecco’s Modified Eagle Medium (DMEM; Thermo Fisher Scientific Inc., MA, USA) supplemented with 10% fetal bovine serum at 37 ℃ in a humidified atmosphere containing 5% CO_2_. Cells between passages 10 and 20 were used for all the experiments. To examine the effects of ANP on renin secretion from As4.1 cells, recombinant mouse ANP (Wuhan Fine Biotech Co., Ltd., Wuhan, China) was added to the culture medium at concentrations of 0.1, 1, and 10 nM. In addition, the NPRA antagonist A71915 (MedChemExpress, NJ, USA) was administered at a concentration of 1 or 5 µM, either alone or in combination with ANP (1 nM). The same treatments were repeated after 12 h, and both the culture supernatants and cells were collected 24 h after the initial treatment.

### RNA extraction and reverse transcription-quantitative PCR

Mouse kidneys were placed in RNAlater Stabilization Solution (Thermo Fisher Scientific) at 4 °C overnight and subsequently stored at −20 °C. Total RNA was extracted from the kidneys or cultured cells using the RNeasy Plus Universal Mini Kit (Qiagen, Hilden, Germany) or the RNeasy Mini Kit (Qiagen). Reverse transcription (RT) was performed using the PrimeScript RT Reagent Kit (Takara Bio Inc., Shiga, Japan) following the manufacturer’s instructions. Quantitative PCR (qPCR) was performed using iTaq Universal SYBR Green Supermix (Bio-Rad Laboratories, Inc., Hercules, CA, USA). The primers used are listed in Table S1. The optimized program was 94 °C for 2 min, followed by 35 cycles of denaturation at 98 °C for 10 s, annealing at 62 °C for 30 s, and extension at 68 °C for 30 s. mRNA expression of target genes was normalized to Gapdh expression. Relative expression levels were determined using the ΔΔCt method and presented as changes compared with the control group.

### RNA-seq

Total RNA used for RNA-seq was extracted from As4.1 cells that were left untreated, treated with ANP alone, or treated with ANP and an NPRA antagonist. cDNA libraries of the RNA samples were subjected to sequencing. Sequencing data were initially quality-checked using FastQC with the FASTQ file reads. Prior to alignment, low-quality reads and adapter sequences were removed using the CutAdapt software. We aligned the FASTQ reads to the GRCm39/ENSEMBL mouse genome using Hisat2. Following alignment, the SAM files were converted to BAM format using Samtools, and the number of mapped reads was counted using HTSeq.

R version 4.4.1 was used for graphical and statistical analyses. DEG analysis was performed using the R package DESeq2 (version 1.46.0) to compare untreated As4.1 cells, cells treated with ANP, and cells treated with ANP and the NPRA antagonist. A volcano plot was generated using ggplot2 in R. Pathway enrichment analysis (GO biological processes) was performed using the ClusterProfiler package.

### Animals

All animals were maintained in a room with controlled temperature and humidity under a 12-hour light/dark cycle. All the animals were handled in accordance with the National Institutes of Health Guide for the Care and Use of Laboratory Animals. This study was approved by the Institutional Animal Care and Ethics Committee of Niigata University (approval number: SA01765). C57BL/6 mice were purchased from Jackson Laboratory Japan, Inc. In the 12 weeks pilot study for mouse model establishment, C57BL/6J mice were used, whereas C57BL/6NJ mice were used for all other 24 weeks experiments.

For experiments investigating afferent arteriolar hypertrophy induced by long-term administration of RAS inhibitors, C57BL/6 mice at 8 weeks of age were divided into three groups: control (vehicle; saline), ARB (valsartan), and ARNI (sacubitril/valsartan). Mice received vehicle (saline), valsartan (30 mg/kg), or sacubitril/valsartan (60 mg/kg) once daily via oral gavage. In the pilot study, mice were divided into groups of n = 12, and 4 mice from each group were sacrificed at 4, 8, and 12 weeks. In the 24 weeks experiment, mice were divided into groups of 16, and all mice were sacrificed at 24 weeks. All mice were fed a low-salt diet (modified to contain 0.1% NaCl; AIN-93G) and had free access to drinking water. Valsartan was purchased from Tokyo Chemical Industry Co., Ltd. (Tokyo, Japan), and sacubitril/valsartan (LCZ696) was obtained from Angene International Ltd. (Nanjing, China).

### Blood pressure measurement

Blood pressure was measured after 24 weeks of treatment. Mice were anesthetized with 1–1.5% isoflurane and maintained at 37.5 °C. Blood pressure was measured under anesthesia using a tail-cuff system (BP-98AL; Softron Ltd., Tokyo, Japan). Three measurements were recorded for each mouse, and the mean value was used as the blood pressure.

### Blood collection

Mice were anesthetized with a three-drug anesthetic mixture (medetomidine, midazolam, and butorphanol), and blood samples were collected via cardiac puncture. Blood was placed into tubes containing EDTA (MiniCollect® II; Greiner Bio-One Co. Ltd., Tokyo, Japan) and into tubes without EDTA. Plasma was separated via centrifugation (1,300 g, 10 min, 4 °C) within 90 min after collection, and serum was obtained after allowing the blood to clot for 30 min followed by centrifugation (1,300 g, 10 min, room temperature) within 2 h. Plasma samples were stored for measurement using ELISA kits. Serum samples were used for a biochemical test panel, which was performed by Oriental Yeast Co. Ltd. (Tokyo, Japan).

### Histological analysis

The mice were sacrificed after 24 weeks, and their kidneys were removed. The excised kidneys were fixed overnight in 10% formalin or Bouin’s solution and embedded in paraffin. Masson’s trichrome and Sirius red staining were performed on 5-µm paraffin sections from kidneys fixed with Bouin’s solution. For Masson’s trichrome staining, sections were deparaffinized in xylene, rehydrated with ethanol, and washed with running water for 5 min. Samples were incubated in Carazzi’s hematoxylin for 30 min, washed under running water for 10 min, and sequentially treated with Ponceau xylidine-acid fuchsin-azophloxine solution for 10 s, 1% acetic acid for 5 min, 2.5% phosphotungstic acid hydrate for 7 min, 1% acetic acid for 5 min, and 1% fast green FCF for 5 min, followed by a final wash in 1% acetic acid for 5 min. The sections were rinsed with running water, dehydrated, cleared, and mounted. For Sirius red staining, the sections were deparaffinized in xylene, rehydrated in ethanol, incubated in resorcin-fuchsin solution for 60 min, rinsed twice in 100% ethanol for 20 s each, and washed with running water for 5 min. The sections were then stained with 3% Picro-Sirius Red solution with gentle agitation for 10 min, dehydrated, cleared, and mounted. The stained sections were visualized under a BZ-X700 microscope (Keyence, Osaka, Japan).

### Immunohistochemistry and Immunofluorescence

For staining for renin and α-smooth muscle actin (α-SMA), kidney sections from paraffin blocks fixed with Bouin’s solution or 10% formalin were deparaffinized in xylenes, rehydrated in 100% ethanol, and washed with phosphate buffered saline (PBS). Sections were then treated with 0.3% hydrogen peroxide in methanol for 30 min. After washing with PBS and blocking with Carbo-Free blocking solution for 30 min at room temperature, the sections were incubated with an anti-renin antibody (ab212197, Abcam, Cambridge, UK) or an anti-α-SMA antibody (A2547, Sigma-Aldrich, USA) at 4 ℃ overnight.

Primary antibodies were used at the following dilutions: anti-renin antibody (1:10,000 for immunohistochemistry [IHC]; 1:1,000 for immunofluorescence [IF]) and anti-α-SMA antibody (1:10,000 for IHC; 1:2,000 for IF). After washing with PBS, the sections were incubated with the secondary antibody. For IHC, goat anti-rabbit IgG (Histofine MAX-PO(R), Nichirei Biosciences Inc., Tokyo, Japan) or goat anti-mouse IgG (Histofine MAX-PO(M), Nichirei Biosciences Inc., Tokyo, Japan) was used as the secondary antibody. For IF, donkey anti-rabbit IgG (H&L) Alexa Fluor 647 and donkey anti-mouse IgG (H+L), highly cross-adsorbed, Alexa Fluor 488 (Thermo Fisher Scientific, Waltham, MA, USA) were used at room temperature for 30 min. For IHC, the sections were developed using the ImmPACT DAB Substrate Kit, Peroxidase (HRP) (SK-4105; Vector Laboratories, Inc., CA, USA). Counterstaining was performed with hematoxylin, followed by rinsing under running water for 30 min. Sections were dehydrated, cleared, and mounted. For IF, the sections were mounted using ProLong Glass Antifade Mountant with NucBlue staining (Invitrogen, Thermo Fisher Scientific, Waltham, MA, USA).

### JG index, thickness of afferent arterioles, and fibrotic area

The JG apparatuses were identified in renin-stained kidney sections in IHC^55^. The JG index was calculated as the ratio of the renin-positive JG apparatuses to the total number of glomeruli and was expressed as a percentage. The thickness of the afferent arterioles was calculated from Masson’s trichrome-stained sections as half the difference between the outer and inner diameters of the afferent arterioles (Figure S3D). The fibrotic area of the kidney was quantified as the red-stained area in the Sirius red-stained sections. The JG index and thickness of afferent arterioles were measured throughout the entire kidney sections, whereas the fibrotic area was quantified in randomly selected cortical regions. All morphometric analyses were performed by investigators who were blinded to the treatment allocation of each sample.

### ELISA

Plasma specimens were obtained from blood after centrifugation at 1,300 g at 4 ℃ for 10 min. The cell culture medium was centrifuged at 1200 × g at room temperature for 10 min and the supernatant was collected. The renin concentration in the plasma and cell culture supernatants was measured using a renin ELISA kit (ELM-Renin1-1; RayBiotech, Norcross, GA, USA). Plasma cystatin C levels were determined using a mouse cystatin C ELISA kit (MSCTC0; R&D Systems, Minneapolis, MN, USA). Plasma ANP levels were measured using an ANP ELISA kit (EIA-ANP-1; RayBiotech, Norcross, GA, USA).

### Single-nucleus RNA sequencing (snRNA-seq)

After 24 weeks of treatment with vehicle, ARB, or ARNI, the mice were anesthetized, and their kidneys were removed. One-quarter of the cortical region from each kidney was snap-frozen and stored for further analysis. Nuclei were isolated from frozen kidney tissue using a Chromium Nuclei Isolation Kit (10x Genomics, Pleasanton, CA, USA) according to the manufacturer’s protocol.

Briefly, three samples from each group were selected, and the frozen kidney tissue was dissociated using a pestle. The dissociated tissues were then mixed according to group and placed on a nuclear isolation column. Nuclei in the flow-through were washed using a debris removal and washing buffer. The concentration of single nuclei was determined using 4′,6-diamidino-2-phenylindole staining. Single nuclei, diluted to a concentration of 1000 nuclei/µL, were captured using the Chromium System (10x Genomics) according to the manufacturer’s instructions. Briefly, the cells were loaded into a Chromium Next GEM Chip G with reagents of Chromium Next GEM single cell 3′ reagent kits v3.1 (10X Genomics). Cell-gel beads were generated in the emulsion and incubated to generate barcoded cDNA. cDNA was purified using Dynabeads (10x Genomics) and amplified using 13 PCR cycles. The cDNA was then enzymatically fragmented, end-repaired, poly-A-tailed, adapter-ligated, and amplified using PCR. The constructed libraries were sequenced on an Illumina Novaseq X Plus platform (150 bp paired-end reads).

The resulting FASTQ file was processed using Cell Ranger v7.0.1 (10x Genomics). The mouse genome (mm10) was used for alignment, and CellRanger count was used for alignment and quantification of gene expression. R version 4.4.1 was used for graphical and statistical analyses. The R package Seurat (version 5.2.1) was used for data analysis, with exclusion criteria for cells with mitochondrial gene percentages exceeding 5% and unique gene counts < 500 or > 4000. After confirming the data quality using DoubletFinder (version 2.0.6), the datasets were integrated using the IntegrateData function implemented in Seurat. Clustering was performed using 12 principal components from the integrated PCA with a resolution of 0.4. UMAP visualization was performed using 12 principal components from the integrated PCA. Cluster identities were assigned using automatic cell-type annotation using ScType. The DEGs between clusters were identified using the Seurat FindMarkers function or MAST (version 1.32.0). Pathway enrichment analysis (GO biological processes) of the DEGs was performed using the ClusterProfiler package. Gene signature scores were calculated using UCell (version 2.10.1), which evaluates predefined gene signatures in single-cell RNA-seq data based on the Mann-Whitney U test^30^. Cell-cell communication analysis was performed using the CellChat package (version 1.6.1)^34^.

### PIC RNA-seq

PIC RNA-seq^35,36^ was performed on mouse kidney tissues. Tissues were fixed in 10% neutral-buffered formalin and embedded in paraffin. Thereafter, 7.5-μm paraffin sections were deparaffinized and permeabilized in TE buffer (pH 8.0) at 70 °C for 1 h, followed by cooling in PBS at 4 °C. Tissue sections were then incubated with an RT reaction mixture comprising a primer mix (0.5 μL of 500 ng/μL NPOM-caged oligo(dT) primer, 0.5 μL of 10 mM dNTPs [NEB], and 5 μL of H₂O) and a first-strand mix (2 μL of 5× First-Strand Buffer, 1 μL of 0.1 M DTT, 0.5 μL of 40 U/μL RNaseOUT, and 0.5 μL of 200 U/μL SuperScript II Reverse Transcriptase; all from Invitrogen) to perform in situ RT. Following in situ RT, sections were blocked with Carbo-Free Blocking Solution for 30 min at room temperature and incubated with an anti-α-SMA antibody (A2547, Sigma-Aldrich; 1:10,000 dilution) for 90 min at room temperature. After washing, sections were incubated with Alexa Fluor 488-conjugated donkey anti-mouse IgG (H + L) and a highly cross-adsorbed secondary antibody (A32766; 1:500; Invitrogen) for 1 h at room temperature. Nuclei were stained with Nuclear Violet LCS1 (17543, AAT Bioquest; 1:500), and the specimens were mounted in SlowFade Diamond (S36972, Thermo Fisher Scientific), followed by coverslipping. Uncaging was performed using light irradiation of α-SMA-positive JG regions.

Illumination was performed using a Leica DM6B fluorescence microscope with a digital mirror device (Polygon1000-G, Mightex) and a 365 nm LED light source (UHP-F-365, Prizmatix). Each region of interest (ROI) was irradiated at 340–380 nm for 3 min. After uncaging, the tissue sections were lysed using proteinase K, and cDNA-mRNA hybrids were collected and purified using the MinElute PCR purification kit (28006, Qiagen). Second-strand cDNA was synthesized using RNase H, DNA polymerase I, and DNA ligase to generate double-stranded cDNA containing a T7 promoter sequence. Double-stranded cDNA was subjected to in vitro transcription using the MEGAscript T7 Transcription Kit (AMB13345; Thermo Fisher Scientific) to generate amplified RNA (aRNA). The aRNA was purified, fragmented, and reverse-transcribed to generate cDNA. The resulting cDNA was amplified using PCR (11 cycles), and library fragments of 250–500 bp were size-selected for library construction using AMPure XP beads through two rounds of purification, and quantified using a Bioanalyzer. Sequencing libraries were subjected to paired-end sequencing on an Illumina NovaSeq 6000 platform, which generated approximately 3 × 106 reads per sample. Read 1 contained sample barcodes and unique molecular identifiers (UMIs), whereas Read 2 contained cDNA sequences.

Raw sequencing data were processed using the PIC analytical pipeline^35,36^. Briefly, barcode and UMI sequences were extracted from Read 1 using UMI-tools (v1.0.0; umi_tools extract -I read1.fastq --read2-in = read2.fastq --bc-pattern = NNNNNNCCCCCC --read2-stdout). Adapter trimming and quality filtering were performed using Trim Galore! (v0.6.6), with the adapter sequence GATCGTCGGACT. Trimmed reads were aligned to the mouse reference genome (GRCm39/mm39) using HISAT2 (v2.1.0) and only uniquely mapped reads were retained for downstream analyses. Gene-level quantification was performed using featureCounts (v1.6.4), and PCR duplicates were removed using UMI-tools (v1.0.0; umi_tools count –method = unique --per-gene --per-cell --gene-tag = XT) to generate UMI-based gene expression matrices. Downstream analyses were conducted in R. Count data were normalized and differential expression analysis was performed using DESeq2 (v1.40.1). Genes with an adjusted p-value < 0.05 were considered differentially expressed. PCA was performed on regularized log-transformed count data using the prcomp function.

### Statistical analysis

Statistical analysis was performed using GraphPad Prism version 11.0.0 (Graphpad Software, La Jolla, CA, USA). The data were analyzed for normal distribution using the Shapiro-Wilk test. Data were considered normally distributed if the *P*-value was >0.05. Normally distributed data were presented as means ± standard deviation. Welch’s t-test or one-way ANOVA with Tukey’s multiple comparison test was used according to the test requirements. P < 0.05 was considered significant. The numbers of replicates and repeats of individual experiments and statistical tests are shown in the figure legends.

## Acknowledgments

We thank Naofumi Imai, Mariko Mikame, and Midori Nagao (Division of Clinical Nephrology and Rheumatology, Kidney Research Center, Niigata University Graduate School of Medicine, Dentistry, and Health Sciences) for their excellent technical assistance.

## References

1. Investigators, H. O. P. E. S. et al. Effects of an Angiotensin-Converting–Enzyme Inhibitor, Ramipril, on Cardiovascular Events in High-Risk Patients. N. Engl. J. Med. 342, 145–153 (2000).

2. Brenner, B. M. et al. Effects of Losartan on Renal and Cardiovascular Outcomes in Patients with Type 2 Diabetes and Nephropathy. N. Engl. J. Med. 345, 861–869 (2001).

3. Dahlöf, B. et al. Cardiovascular morbidity and mortality in the Losartan Intervention For Endpoint reduction in hypertension study (LIFE): a randomised trial against atenolol. Lancet 359, 995–1003 (2002).

4. Whelton, P. K. et al. 2017 ACC/AHA/AAPA/ABC/ACPM/AGS/APhA/ASH/ASPC/NMA/PCNA Guideline for the Prevention, Detection, Evaluation, and Management of High Blood Pressure in Adults: A Report of the American College of Cardiology/American Heart Association Task Force on Clinical Practice Guidelines. Hypertension 71, e13–e115 (2018).

5. Riet, L. te, Esch, J. H. M. van, Roks, A. J. M., Meiracker, A. H. van den & Danser, A. H. J. Hypertension: renin-angiotensin-aldosterone system alterations. Circ. Res. 116, 960–975 (2015).

6. Gomez, R. A. & Sequeira-Lopez, M. L. S. Renin cells in homeostasis, regeneration and immune defence mechanisms. Nat. Rev. Nephrol. 14, 231–245 (2018).

7. Watanabe, H. et al. Renin Cell Baroreceptor, a Nuclear Mechanotransducer Central for Homeostasis. Circ Res 129, 262–276 (2021).

8. Sequeira-Lopez, M. L. S., Pentz, E. S., Nomasa, T., Smithies, O. & Gomez, R. A. Renin cells are precursors for multiple cell types that switch to the renin phenotype when homeostasis is threatened. Dev Cell 6, 719–728 (2004).

9. Guessoum, O., Martini, A. de G., Sequeira-Lopez, M. L. S. & Gomez, R. A. Deciphering the Identity of Renin Cells in Health and Disease. Trends Mol Med 27, 280–292 (2020).

10. Yamaguchi, H., Gomez, R. A. & Sequeira-Lopez, M. L. S. Renin Cells, From Vascular Development to Blood Pressure Sensing. Hypertension 80, 1580–1589 (2023).

11. Oka, M., Medrano, S., Sequeira-Lόpez, M. L. S. & Gómez, R. A. Chronic Stimulation of Renin Cells Leads to Vascular Pathology. Hypertension 70, 119–128 (2018).

12. Watanabe, H., et al. Inhibition of the renin-angiotensin system causes concentric hypertrophy of renal arterioles in mice and humans. JCI Insight 6, e154337 (2021).

13. Nakanishi, K., Nagai, Y., Akimoto, T. & Yamanaka, N. Changes in renal vessels associated with long-term administration of angiotensin converting enzyme inhibitor in Zucker fatty rats. J. Smooth Muscle Res. 53, 20–30 (2017).

14. Nagai, Y., Nakanishi, K., Akimoto, T. & Yamanaka, N. Proliferative changes of renal arteriolar walls induced by administration of angiotensin II receptor blocker are frequent in juvenile rats. J. Renin-Angiotensin-Aldosterone Syst. 15, 440–449 (2014).

15. Yamaguchi, M. et al. Transformation of the Kidney into a Pathological Neuro-Immune-Endocrine Organ. Circ. Res. (2024) doi:10.1161/circresaha.124.325305.

16. Nagai, Y., et al. A Study of Morphological Changes in Renal Afferent Arterioles Induced by Angiotensin II Type 1 Receptor Blockers in Hypertensive Patients. Kidney Blood Press Res 45, 194–208 (2020).

17. Almeida, L. F. de et al. Balancing Blood Pressure Targets and Kidney Health: The Potential Risk of Kidney Disease Due to Concentric Arterial and Arteriolar Hypertrophy From Renin-Angiotensin-Aldosterone Inhibitors. Hypertension (2026) doi:10.1161/hypertensionaha.125.26125.

18. Maack, T. Role of atrial natriuretic factor in volume control. Kidney Int. 49, 1732–1737 (1996).

19. McFarlane, S. I., Winer, N. & Sowers, J. R. Role of the Natriuretic Peptide System in Cardiorenal Protection. Arch. Intern. Med. 163, 2696–2704 (2003).

20. Volpe, M. Natriuretic peptides and cardio-renal disease. Int. J. Cardiol. 176, 630–639 (2014).

21. Devlin, A. M. & Leckie, B. J. Atrial natriuretic peptide suppresses isoprenaline and dibutyryl cyclic adenosine monophosphate-induced cell growth in cultured renin-secreting human nephroblastoma cells Comparison with forskolin-induced renin secretion. J. Hypertens. 10, 445–450 (1992).

22. Kurtz, A., Bruna, R. D., Pfeilschifter, J., Taugner, R. & Bauer, C. Atrial natriuretic peptide inhibits renin release from juxtaglomerular cells by a cGMP-mediated process. Proc. Natl. Acad. Sci. 83, 4769–4773 (1986).

23. Volpe, M., Carnovali, M. & Mastromarino, V. The natriuretic peptides system in the pathophysiology of heart failure: from molecular basis to treatment. Clin. Sci. (Lond., Engl. : 1979) 130, 57–77 (2016).

24. Singh, J. S. S. et al. Sacubitril/valsartan: beyond natriuretic peptides. Heart 103, 1569 (2017).

25. Haynes, R. et al. Effects of Sacubitril/Valsartan Versus Irbesartan in Patients With Chronic Kidney Disease. Circulation 138, 1505–1514 (2018).

26. Barbosa, C. V., Lang, H., Melk, A. & Schmidt, B. M. W. Renal events in patients receiving neprilysin inhibitors: a systematic review and meta-analysis. Nephrol. Dial. Transplant. 37, 2418–2428 (2022).

27. McMurray, J. J. V. et al. Angiotensin–Neprilysin Inhibition versus Enalapril in Heart Failure. N. Engl. J. Med. 371, 993–1004 (2014).

28. Causland, F. R. M. et al. Angiotensin-Neprilysin Inhibition and Renal Outcomes in Heart Failure With Preserved Ejection Fraction. Circulation 142, 1236–1245 (2020).

29. Damman, K. et al. Renal Effects and Associated Outcomes During Angiotensin-Neprilysin Inhibition in Heart Failure. JACC: Hear. Fail. 6, 489–498 (2018).

30. Andreatta, M. & Carmona, S. J. UCell: Robust and scalable single-cell gene signature scoring. Comput. Struct. Biotechnol. J. 19, 3796–3798 (2021).

31. Neri, D. et al. Distinct sympathetic projections to brown fat regulate thermogenesis and glucose tolerance. Nat. Metab. 1–14 (2026) doi:10.1038/s42255-025-01429-0.

32. Minnone, G., Benedetti, F. D. & Bracci-Laudiero, L. NGF and Its Receptors in the Regulation of Inflammatory Response. Int. J. Mol. Sci. 18, 1028 (2017).

33. Gauthier, M. M., Hayoz, S. & Banek, C. T. Neuroimmune interplay in kidney health and disease: Role of renal nerves. Auton. Neurosci. 250, 103133 (2023).

34. Jin, S. et al. Inference and analysis of cell-cell communication using CellChat. Nat. Commun. 12, 1088 (2021).

35. Honda, M. et al. High-depth spatial transcriptome analysis by photo-isolation chemistry. Nat. Commun. 12, 4416 (2021).

36. Honda, M. et al. Photo-isolation chemistry for high-resolution and deep spatial transcriptome with mouse tissue sections. STAR Protoc. 3, 101346 (2022).

37. Group, K. D. I. G. O. (KDIGO) B. P. W., et al. KDIGO 2021 Clinical Practice Guideline for the Management of Blood Pressure in Chronic Kidney Disease. Kidney Int. 99, S1–S87 (2021).

38. Lewis, E. J. et al. Renoprotective Effect of the Angiotensin-Receptor Antagonist Irbesartan in Patients with Nephropathy Due to Type 2 Diabetes. N. Engl. J. Med. 345, 851–860 (2001).

39. Berg, A. C., Chernavvsky-Sequeira, C., Lindsey, J., Gomez, R. A. & Sequeira-Lopez, M. L. S. Pericytes synthesize renin. World J. Nephrol. 2, 11 (2013).

40. Meyrier, A. Nephrosclerosis: update on a centenarian. Nephrol. Dial. Transplant. 30, 1833–1841 (2015).

41. Kopp, J. B. Rethinking hypertensive kidney disease. Curr. Opin. Nephrol. Hypertens. 22, 266–272 (2013).

42. Kohagura, K., Zamami, R., Oshiro, N., Shinzato, Y. & Uesugi, N. Heterogeneous afferent arteriolopathy: a key concept for understanding blood pressure–dependent renal damage. Hypertens. Res. 1–14 (2024) doi:10.1038/s41440-024-01916-z.

43. Pentz, E. S., Moyano, M. A., Thornhill, B. A., Lopez, M. L. S. S. & Gomez, R. A. Ablation of renin-expressing juxtaglomerular cells results in a distinct kidney phenotype. Am. J. Physiol.-Regul., Integr. Comp. Physiol. 286, R474–R483 (2004).

44. Shi, S.-J., Nguyen, H. T., Sharma, G. D., Navar, L. G. & Pandey, K. N. Genetic disruption of atrial natriuretic peptide receptor-A alters renin and angiotensin II levels. Am. J. Physiol.-Ren. Physiol. 281, F665–F673 (2001).

45. Pandey, K. N. & Vellaichamy, E. Regulation of cardiac angiotensin-converting enzyme and angiotensin AT1 receptor gene expression in Npr1 gene-disrupted mice. Clin. Exp. Pharmacol. Physiol. 37, e70–e77 (2010).

46. Volpe, M., Gallo, G. & Rubattu, S. Endocrine functions of the heart: from bench to bedside. Eur. Hear. J. 44, 643–655 (2022).

47. Khurana, M. L., Mani, I., Kumar, P., Ramasamy, C. & Pandey, K. N. Ligand-Dependent Downregulation of Guanylyl Cyclase/Natriuretic Peptide Receptor-A: Role of miR-128 and miR-195. Int. J. Mol. Sci. 23, 13381 (2022).

48. Kumar, R., Geldern, T. W. von, Calle, R. A. & Pandey, K. N. Stimulation of atrial natriuretic peptide receptor/guanylyl cyclase-A signaling pathway antagonizes the activation of protein kinase C-α in murine Leydig cells. Biochim. Biophys. Acta (BBA) - Mol. Cell Res. 1356, 221–228 (1997).

49. Potter, L. R. Natriuretic peptide metabolism, clearance and degradation. FEBS J. 278, 1808–1817 (2011).

50. Williams, B. et al. Effects of Sacubitril/Valsartan Versus Olmesartan on Central Hemodynamics in the Elderly With Systolic Hypertension. Hypertension 69, 411–420 (2017).

51. Han, Y., Zhou, Y., Na, J., Li, F. & Sun, Y. Efficacy and Safety Comparative of Sacubitril/Valsartan vs. Olmesartan in the Treatment of hypertension: A Meta-analysis of RCTs. Am. J. Hypertens. 36, 643–650 (2023).

52. Causland, F. R. M. et al. Angiotensin–neprilysin inhibition and renal outcomes across the spectrum of ejection fraction in heart failure. Eur. J. Hear. Fail. 24, 1591–1598 (2022).

53. D’Elia, E. et al. Neprilysin inhibition in heart failure: mechanisms and substrates beyond modulating natriuretic peptides. Eur. J. Hear. Fail. 19, 710–717 (2017).

54. Sigmund, C. D. et al. Isolation and characterization of renin-expressing cell lines from transgenic mice containing a renin-promoter viral oncogene fusion construct. J. Biol. Chem. 265, 19916–22 (1990).

55. Yamaguchi, H. et al. Krüppel-like factor 2 regulates renin expression in mature juxtaglomerular cells. Am. J. Physiol.-Ren. Physiol. 330, F410–F421 (2026).

