## Supplemental Material for "Natriuretic Peptide Augmentation Attenuates Renin Cell Hyperactivation and Afferent Arteriolar Hypertrophy During Long-Term Renin-Angiotensin System Inhibition"

**Table S1. Primers for reverse transcription-quantitative PCR.**

| Gene | Forward sequence (5' to 3') | Reverse sequence (5' to 3') |
|---|---|---|
| *Ren1* | ACATGACCAGGCTCAGTGCTGA | TACCGATGCCAATCTCGCCGTA |
| *Havcr1* | CTGGAATGGCACTGTGACATCC | GCAGATGCCAACATAGAAGCCC |
| *Lcn2* | ATGTCACCTCCATCCTGGTCAG | GCCACTTGCACATTGTAGCTCTG |
| *Col1a1* | GAGCGGAGAGTACTGGATCG | TACTCGAACGGGAATCCATC |
| *Col3a1* | TCCTAACCAAGGCTGCAAGATGGA | ACCAGAATCTGTCCACCAGTGCTT |
| *Acta2* | ATTGTGCTGGACTCTGGAGATGGT | TGATGTCACGGACAATCTCACGCT |
| *Ngf* | GTTTTGCCAAGGACGCAGCTTTC | GTTCTGCCTGTACGCCGATCAA |
| *Gapdh* | CATCACTGCCACCCAGAAGACTG | ATGCCAGTGAGCTTCCCGTTCAG |

**Figure S1**

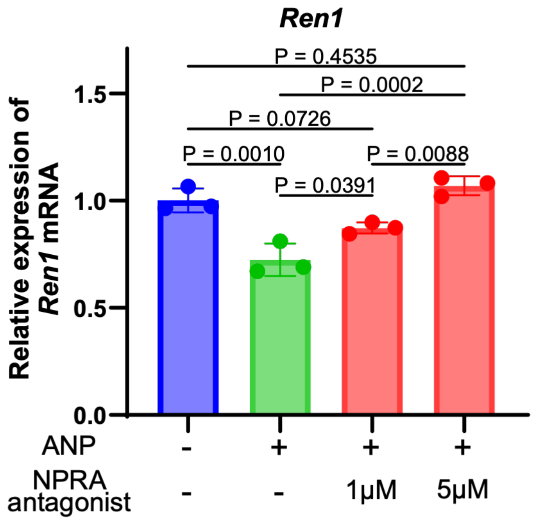


**Figure S1. Effects of ANP and natriuretic peptide receptor A antagonism on *Ren1* expression in As4.1 cells.**

Relative *Ren1* mRNA expression in As4.1 cells 24 hours after treatment with vehicle, ANP (1 nM) alone, or ANP (1nM) in combination with the natriuretic peptide receptor A antagonist A71915 (1 or 5 μM). Data are shown as mean ± SD (n=3 per group in the representative experiment). Statistical significance was assessed by one-way ANOVA followed by Tukey's multiple-comparisons test. Data are representative of three independent experiments.

**Figure S2**

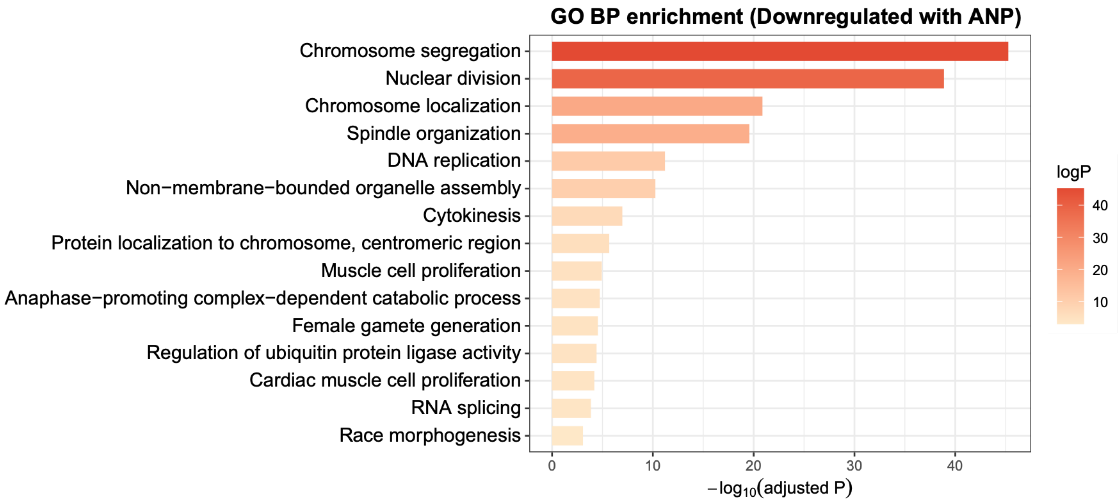


**Figure S2. Gene Ontology enrichment analysis of ANP-downregulated genes in As4.1 cells.**

Gene Ontology (GO) Biological Process enrichment analysis of genes downregulated following ANP treatment in As4.1 cells.

**Figure S3**

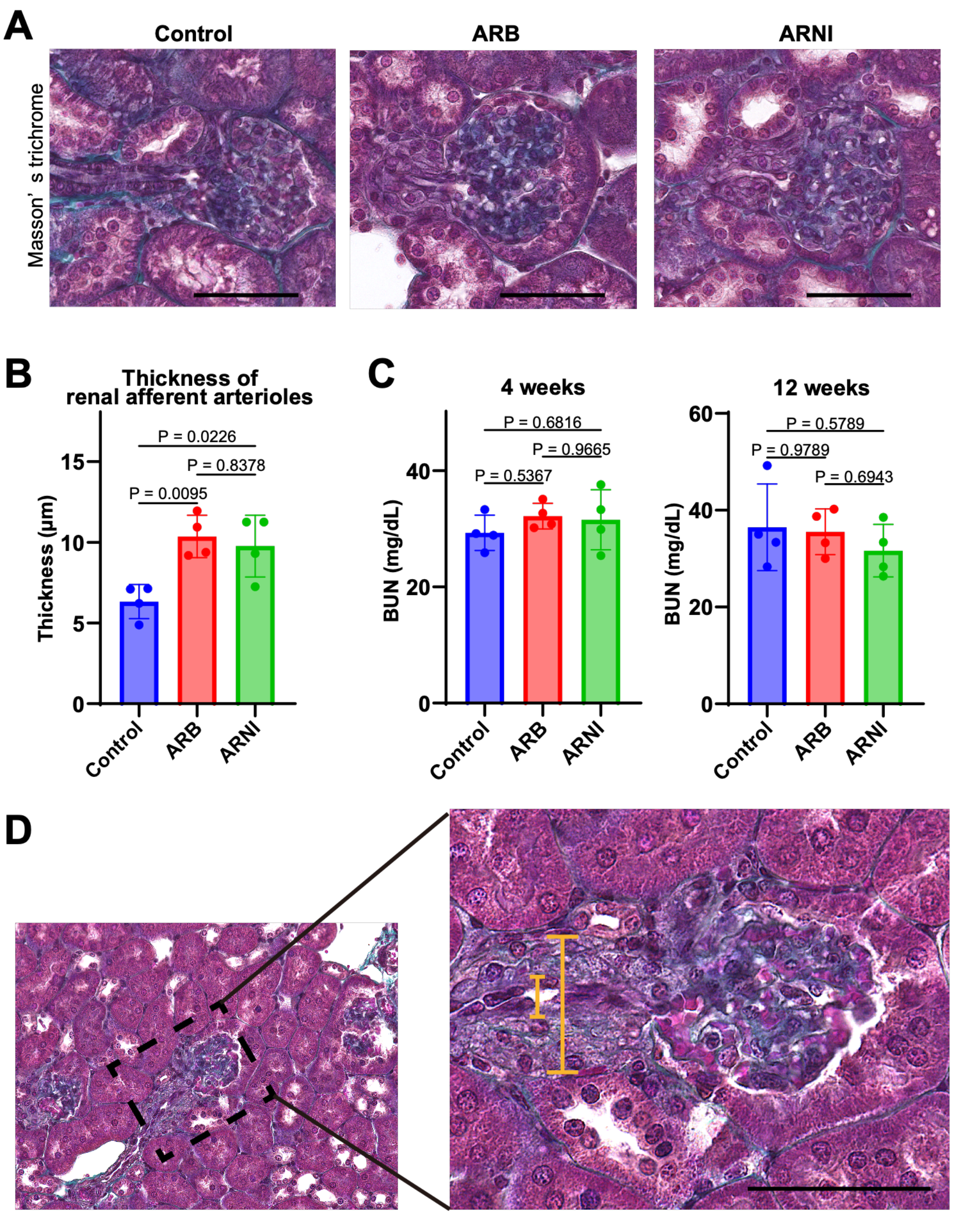


**Figure S3. Pilot study of mice treated with RAS inhibitors for 4–12 weeks.**

**A**, Representative Masson's trichrome-stained kidney sections after 12 weeks of treatment. Scale bar, 50 μm. **B**, Quantification of afferent arteriolar wall thickness after 12 weeks of treatment. Afferent arteriolar wall thickness was significantly increased in both the ARB and ARNI groups compared with the control group. n=4 per group; one-way ANOVA followed by Tukey's multiple-comparisons test. **C**, Serum blood urea nitrogen (BUN) levels measured at 4 and 12 weeks after treatment initiation. No significant differences were observed among the groups at either time point. n=4 per group at each time point; one-way ANOVA followed by Tukey's multiple-comparison test. **D**, Schematic illustrating the measurement of afferent arteriolar wall thickness in Masson's trichrome-stained sections. Wall thickness was calculated as half the difference between the outer and inner diameters of the afferent arteriole.

### Figure S4

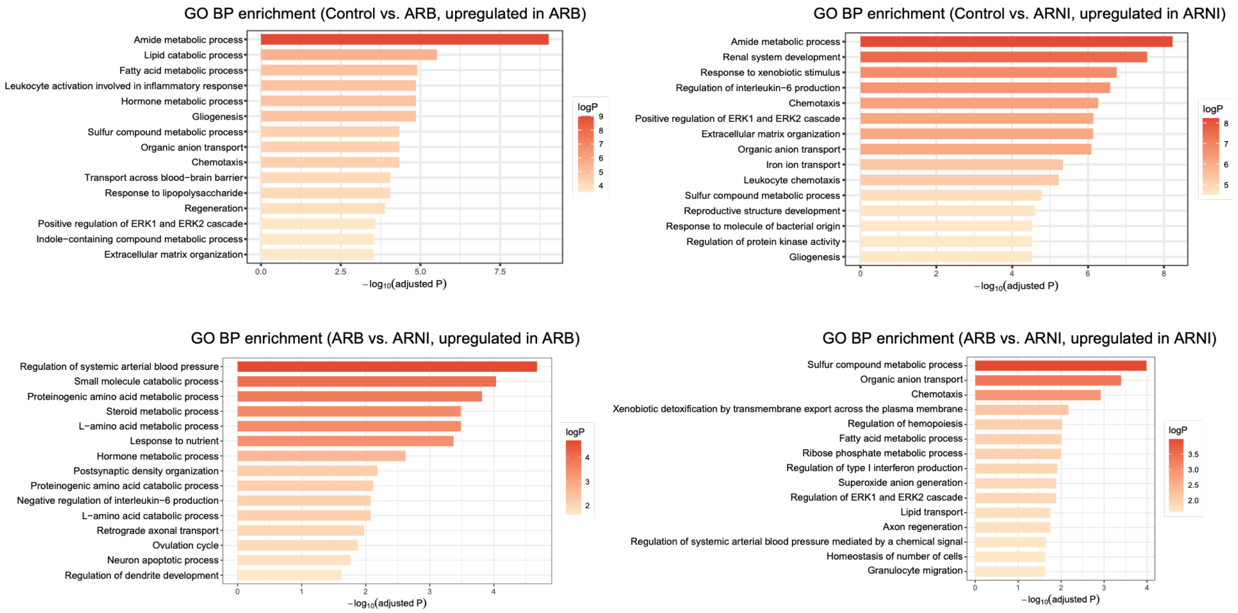


**Figure S4. Gene Ontology enrichment analysis of proximal tubular transcriptional responses during long-term RAS inhibition.**

Gene Ontology (GO) Biological Process enrichment analysis of differentially expressed genes in proximal tubule clusters among the control, ARB, and ARNI groups. Enrichment results are shown for genes upregulated in the ARB or ARNI groups compared with the control group, as well as for genes differentially expressed between the ARB and ARNI groups.

**Figure S5**

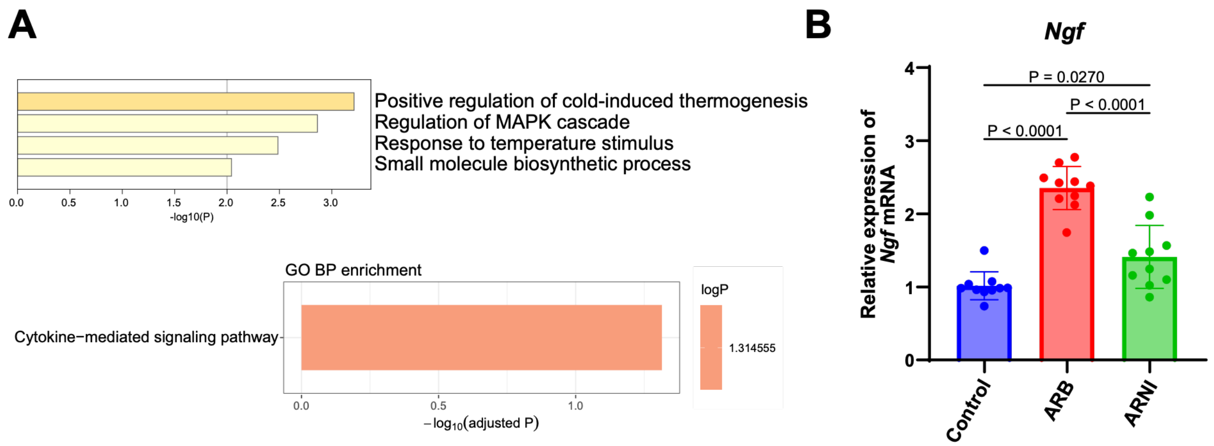


**Figure S5. Mesenchymal transcriptional programs and renal *Ngf* expression during long-term RAS inhibition.**

**A**, Gene Ontology (GO) Biological Process enrichment analysis of differentially expressed genes in the mesenchymal cell cluster between the ARB and ARNI groups. Enriched terms associated with genes downregulated in the ARNI group relative to the ARB group are shown. **B**, Relative *Ngf* mRNA expression in whole-kidney samples. Long-term ARB treatment significantly increased *Ngf* expression, whereas this increase was attenuated in the ARNI group. n=10 per group; one-way ANOVA followed by Tukey’s multiple-comparison test.
